# Refined measurement of SecA-driven protein transport reveals indirect coupling to ATP turnover

**DOI:** 10.1101/2020.05.08.084160

**Authors:** William J. Allen, Daniel W. Watkins, Mark S. Dillingham, Ian Collinson

**Affiliations:** University of Bristol

## Abstract

The universally conserved Sec system is the primary method cells utilise to transport proteins across membranes. Until recently, measuring the activity – a prerequisite for understanding how biological systems works – has been limited to discontinuous protein transport assays with poor time resolution, or used as reporters large, non-natural tags that interfere with the process. The development of an assay based on a split super-bright luciferase (NanoLuc) changed this. Here, we exploit this technology to unpick the steps that constitute post-translational transport in bacteria. Under the conditions deployed, transport of the model pre-protein substrate proSpy occurs at 200 amino acids per minute with the data best fit by a series of large, ∼30 amino acid, steps each coupled to many (100s) ATP hydrolysis events. Prior to that, there is no evidence for a distinct, rate-limiting initiation event. Kinetic modelling suggests that SecA-driven transport activity is facilitated by the substrate (polypeptide) concentration gradient – in keeping with classical membrane transporters. Furthermore, the features we describe are consistent with a non-deterministic motor mechanism, such as a Brownian ratchet.

## Introduction

To transport proteins from one side of a lipid bilayer to the other, cells employ specialised, membrane-embedded molecular motors. These recognise proteins for transport, then use energy from ATP binding and hydrolysis and/or the proton-motive force (PMF) to transfer them through a polypeptide conducting channel in the membrane – usually threading them through in an unfolded state. Probably the best studied protein transporter is the Sec system of *Escherichia coli*, which handles almost all proteins destined for the cell envelope and beyond. In its post-translational mode – used for exporting periplasmic, outer membrane and extra-cellular proteins – the cytosolic ATPase SecA binds pre-proteins with a cleavable N-terminal signal sequence (SS; proteins with the SS present are termed pre-proteins), and translocates them through the membrane-embedded heterotrimeric core-complex SecYEG.

In many ways, the bacterial Sec system is quite well characterised: several structures are available of the channel complex and motor ATPase SecA, alone and associated (Berg et al., 2004; Hunt et al., 2002; Zimmer et al., 2008); the pathway the pre-protein takes and how various domains move have been mapped extensively using biochemical, biophysical and computational approaches (Bauer and Rapoport, 2009; Cannon et al., 2005; Corey et al., 2019; Gold et al., 2012; Li et al., 2016); and the ATPase activity of SecA and its regulation have been subject to detailed dissection (Allen et al., 2016; Economou et al., 1995; Gold et al., 2007; Lill et al., 1989; Robson et al., 2009). Yet despite this, the Sec field has yet to agree on an answer to the question: how is ATP hydrolysis actually coupled to protein transport?

Perhaps the biggest barrier to elucidating ‘the’ mechanism of Sec is the huge variability of peptide sequences in terms of size, shape and chemistry. Unlike motors that run along DNA or RNA – which have a repeating sugar-phosphate backbone to grip onto – or those that move along predictably organised cytoskeletal helical filaments, protein transporters must by turns recognise hydrophobic and hydrophilic regions, small and bulky residues, with both positive and negative charge and varying amounts of secondary structure. Therefore, no single set of domain movements is likely to work for every part of every pre-protein. Instead, current models for Sec are not purely deterministic: they allow at least some measure of pre-protein diffusion through the channel. We have previously proposed a pure ratcheted diffusion model (Ahdash et al., 2019; Allen et al., 2016; Corey et al., 2019), while others have proposed a hybrid ‘push and slide’ model (Bauer et al., 2014).

Very precise measurements of protein transport are required to distinguish between these types of mechanism, such as those produced by the recently published NanoLuc transport assay (Pereira et al., 2019). Here we extend the use of the NanoLuc assay to reveal the elementary steps of the protein transport mechanism, using the model pre-protein pro-spheroplast protein Y (pSpy). The results reveal a highly non-deterministic transport model, with a small apparent number of steps each of which requires hundreds of ATP turnovers to resolve *in vitro*. The overall rate of transport is about 200 amino acids per minute, dependent on the pre-protein concentration gradient across the membrane but not limited by a distinct initiation step.

## Results

### Using NanoLuc to dissect translocation kinetics

To interrogate the kinetics of protein transport in sufficient detail to glean mechanistic information we used the recently developed NanoLuc system (Pereira et al., 2019). In essence, NanoLuc luciferase missing a single β-strand (11S) is encapsulated within proteo-liposomes (PLs) incorporating the Sec machinery, while a high affinity version of the missing β-strand (Pep86) is fused to a translocation substrate. These are then mixed together in the presence of the luciferase substrate furimazine and an ATP regeneration system, allowed to equilibrate, and the reaction started by the addition of ATP. As pre-protein is transported into the PLs, Pep86 complements 11S producing a luminescent signal. An example import curve, with background subtracted (Fig. S1a; Pereira et al., 2019) is shown in Fig. 1a. It can be fitted, as a fairly good approximation, to a simple delay phase (**lag**) followed by a single exponential with an apparent rate constant (**λ**) and amplitude (**A**; Fig. 1a).

**Figure 1:**
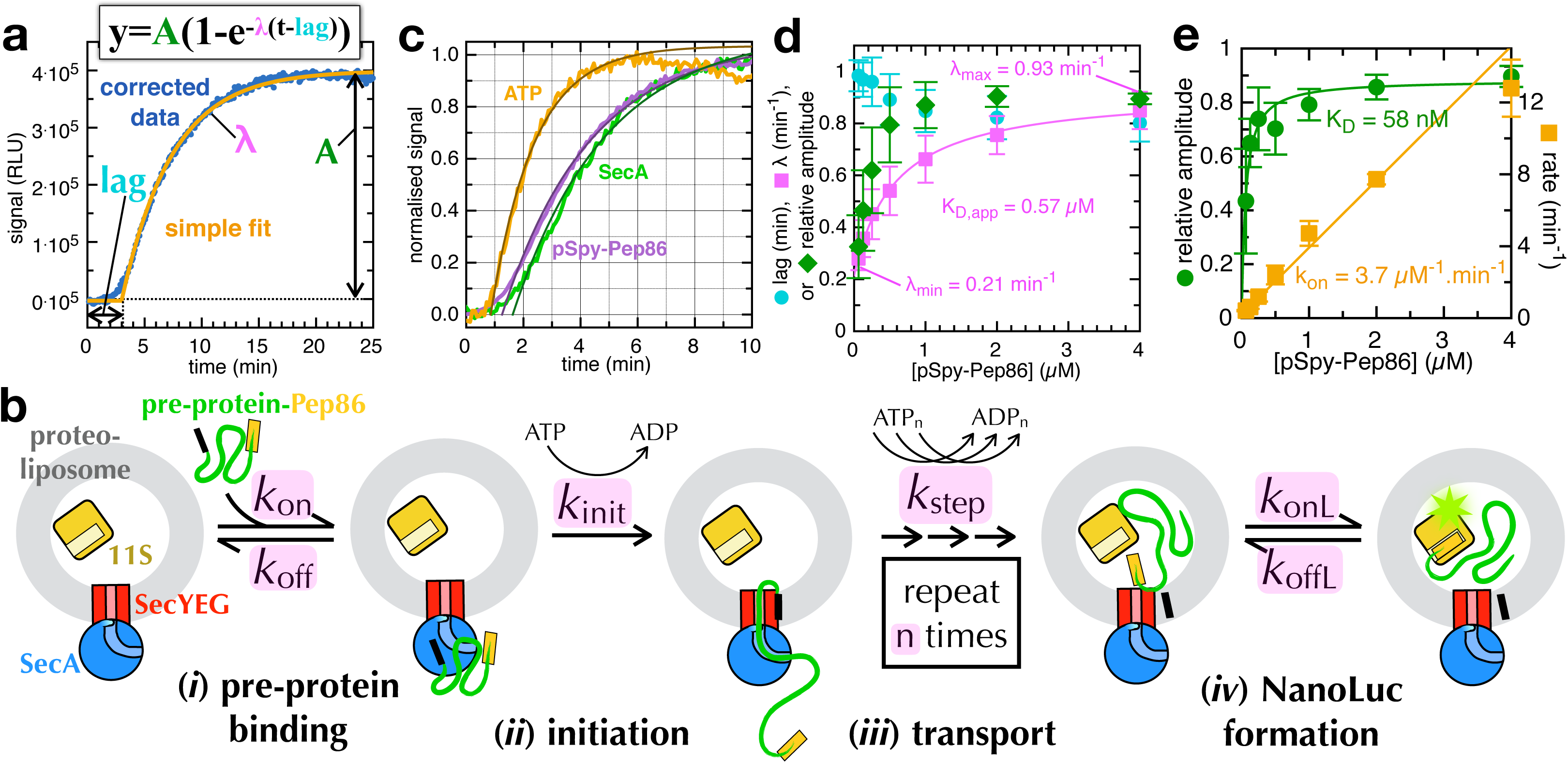
The NanoLuc transport assay. **a)** An example NanoLuc transport curve, with simple fitting. **b)** The minimal model used to describe pre-protein import in the NanoLuc assay. **c)** Transport initiated by the addition ATP (orange), pSpy-Pep86 (purple) or SecA (green). Lines represent best fit to the single exponential + lag model. **d)** Fitted λ (pink squares), amplitude (green diamonds) and lag (cyan circles) as a function of pSpy-Pep86 concentration. Error bars represent average and SEM from 4 repeats. λ is fitted to a weak binding equation. **e)** Secondary data from a titration of pSpy-Pep86 against 200 pM 11S in solution. Fits are to a weak binding equation for amplitude (green circles) and a straight line for rate (orange squares).

The observed kinetics are characteristic of “n-step sequential” translocation mechanisms such as those that have been applied to the analysis of DNA helicase motors (Fig. 1b; Ali and Lohman, 1997; Lucius et al., 2003; McClelland et al., 2005) and the lag-exponential fit can be used to extract semi-quantitative information from the data. The amplitude (**A**) corresponds to the amount of NanoLuc that is formed when the reaction reaches completion. The **lag** before any signal occurs is the result of the accumulation of transport intermediates, because translocation of the protein involves multiple consecutive steps with comparable rates (Lucius et al., 2003). It corresponds to the sum of the time constants (i.e. ∑1/*k*_all_) for all steps prior to the one that yields the signal (in our case the final NanoLuc formation step; McClelland et al., 2005). The **λ** factor is complex and has a less obvious physical meaning with respect to the stepping mechanism. It contains information related to rate limiting steps in the overall pathway, as well as both the translocation step size and the static disorder in the stepping rate; we return to this point later.

Taking the observed kinetics into account alongside prior knowledge of the sequence of events leading to protein secretion, a minimal reaction scheme for NanoLuc-monitored protein import can be devised, which we will use as an initial framework for the analysis (Fig. 1b). Firstly, the pre-protein substrate must be recognised by the SecYEG-SecA complex (step *i*; on- and off-rates *k*_on_ and *k*_off_, respectively). Note that in our setup this step starts at equilibrium, as it does not require additional energy input. Recognition is followed by an ATP-dependent initiation step (*ii*, rate *k*_init_), wherein the signal sequence unlocks the channel and primes it for transport (Corey et al., 2016; Fessl et al., 2018; Gouridis et al., 2009; Hizlan et al., 2012; Li et al., 2016). Transport itself (step *iii*) is driven by a number (n) of ATP-dependent steps, each of which has the rate *k*_step_. In a physiological context, these would presumably be assisted by the PMF (Brundage et al., 1990; Schiebel et al., 1991). Finally, once the Pep86 at the C-terminus of the protein has been transported into the PL, it must associate with 11S to form mature NanoLuc (step *iv*, with on- and off-rates *k*_onL_ and *k*_offL_, respectively).

All these four steps must occur in order for us to measure a transport signal, but this does not necessarily mean they all contribute appreciably to the kinetics. For example, step (*ii*) is included in the model because there is ample experimental evidence that it is important for recognising secretory substrates (Fessl et al., 2018; Gouridis et al., 2009), but it might be too fast to affect the shape of the transport curves. Furthermore, even this relatively simple model makes some basic assumptions, e.g. that the initiated complex never dissociates (infinite processivity), discussed below.

### Establishing a minimal model for transport

To explore how the parameters in the above model are related to the observed data, we carried out some control experiments related to step *i* and *iv*. First, we compared reactions initiated by the addition of ATP (Fig. 1c, orange line) with those initiated by pSpy-Pep86 or SecA (Fig. 1c, purple and green lines, respectively). The only difference between these runs is whether the step (*i*) is at equilibrium when the reaction starts (ATP), or if the SecYEG-SecA-pre-protein complex must form first (SecA or pSpy-Pep86). The difference in lag between the pre-equilibrated reaction (lag_ATP_ = 0.85 min) and the two reactions where step *i* must also take place (lag_pSpy_ = 1.25 min; lag_SecA_ = 1.66 min) will therefore equal the *k*^−1^ of assembly of the pre-initiation complex. For 2 μM pSpy, this is 2.5 min^−1^ (1÷(1.25-0.85)), or 1.25 μM^−1^.min^−1^; while with 1 μM SecA it is 1.23 min^−1^ (1 ÷(1.66-0.85)) – also ∼1.25 μM^−1^.min^−1^. The big difference between the reaction curves also suggests that most transport signal derives from pre-initiated complex – i.e. transport is largely single turnover under the conditions used here. This is also supported by experiments where excess pSpy without Pep86 is added along with the ATP (Fig. S1b-c). However, as there is nothing to stop fresh cycles of transport starting even after ATP is added, this presumably reflects the amount of pre-formed complex and will not be the case under all reaction conditions. Because we are most interested in extracting *k*_init_, *k*_step_ and n, all subsequent experiments were initiated using ATP.

We next investigated the effect of titrating pSpy-Pep86 concentration on transport. Import signal fits well to the simple exponential + lag fit (example raw data are shown in Fig. S1d, and normalised in Fig. S1e). The best fit parameters plotted as a function of pSpy concentration (Fig. 1d; error bars are the SEM from four repeats) show that amplitude and λ (Fig. 1d, green and pink respectively) are both strongly affected by pre-protein concentration, while lag (Fig. 1d, cyan) is affected little if at all. Assuming transport takes place according to the model in Fig. 1b (and at least some pSpy-Pep86 is pre-bound), lag should correspond to:

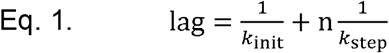

The observation that lag is not pre-protein concentration dependent is therefore expected, as none of the parameters that define it are either. The plot of λ as a function of pSpy-Pep86 concentration, meanwhile, fits fairly well to a weak binding equation (Fig. 1d, pink squares) giving an apparent K_D_ (0.57 μM) in reasonable agreement with the previously determined affinity of pSpy for SecA (0.2 μM; (Chatzi et al., 2017)).

The fact that signal amplitude depends on pre-protein concentration was surprising to us: although lower pre-protein concentrations mean less pre-formed SecYEG-SecA-pre-protein complex, there is no obvious reason why transport should not continue to initiate even after the addition of ATP. PLs are diluted approximately 8000-fold (w/v) into reaction buffer for the transport experiment. So for an internal concentration of 11S of 40 μM (the maximum used here), only 5 nM pre-protein needs to be imported to completely saturate all the encapsulated 11S. The concentration of pSpy-Pep86 is therefore essentially unaffected by transport; the reaction is reaching completion in a manner that is dependent on pSpy-Pep86 concentration, but is not due to it running out – discussed below.

It should be noted that signal amplitude (normalised to its maximum value within each repeat) against [pSpy-Pep86] produces a graph with extremely large error bars (Fig. 1d, green diamonds). Presumably, these reflect that fact that signal amplitude is very sensitive to multiple different experimental parameters – especially 11S concentration within the PLs, which is highly variable between batches. To mitigate against this, all subsequent experiments where comparisons are made between different experimental conditions are performed using the same batch of PLs, and in parallel on a plate reader where possible. It should also be mentioned that when the value for λ is very low it becomes hard to determine the x-intercept precisely; thus lags determined from transport reactions that reach completion quickly are more reliable.

Finally, we measured NanoLuc formation (step *iv*) in solution (no membranes present) by titrating pSpy-Pep86 against a fixed concentration of 11S. The rate of formation is approximately linear up to 4 μM (above which it becomes too fast to resolve on the plate reader) with a slope (*k*_onL_) of 3.7 μM^−1^.min^−1^ (Fig. 1e, orange squares). The fitted K_D_ for the interaction (K_D,L_) is 58 ± 24 nM (Fig. 1e, green circles; error derived from the fit), which means that for 11S concentrations used here (generally 10-40 μM inside the PL), NanoLuc formation should always be completely saturated and very fast relative to transport (see e.g. Fig. 1c-d, and below). Consistent with this, we have previously shown that the concentration of 11S inside the vesicles has no effect on transport kinetics of a different model substrate (proOmpA) at concentrations above 1 μM (Pereira et al., 2019).

### Determination of the initiation and transport steps using tandem pSpys

To investigate the protein transport parameters *k*_init_, *k*_step_ and n, we next designed a series of four nearly identical 4x tandem pSpy-Pep86 variants (pSpy_4x_; Fig. 2a, Fig. S2). In each substrate, three of the Pep86 sequences are scrambled so they retain the same amino acid composition but give a vastly reduced signal upon transport (Fig. S3a; ‘D’ (for dark) in Fig. 2a). The fourth is left as active Pep86 (’L’ (for light) in Fig. 2a). Thus, the resulting proteins are identical save for the length of substrate that must be translocated before the functional Pep86 becomes accessible. After confirming that all four bind rapidly and with high affinity to 11S (Fig. S3b-e), we carried out transport experiments initiated with saturating ATP (Fig. 2b). We observed that, as the position of the active Pep86 moves later (from LDDD to DDDL, Fig. 2b) all three parameters in the lag-exponential fit are affected in a systematic manner: the lag increases with the length of substrate transported, while λ and amplitude both decrease.

**Figure 2:**
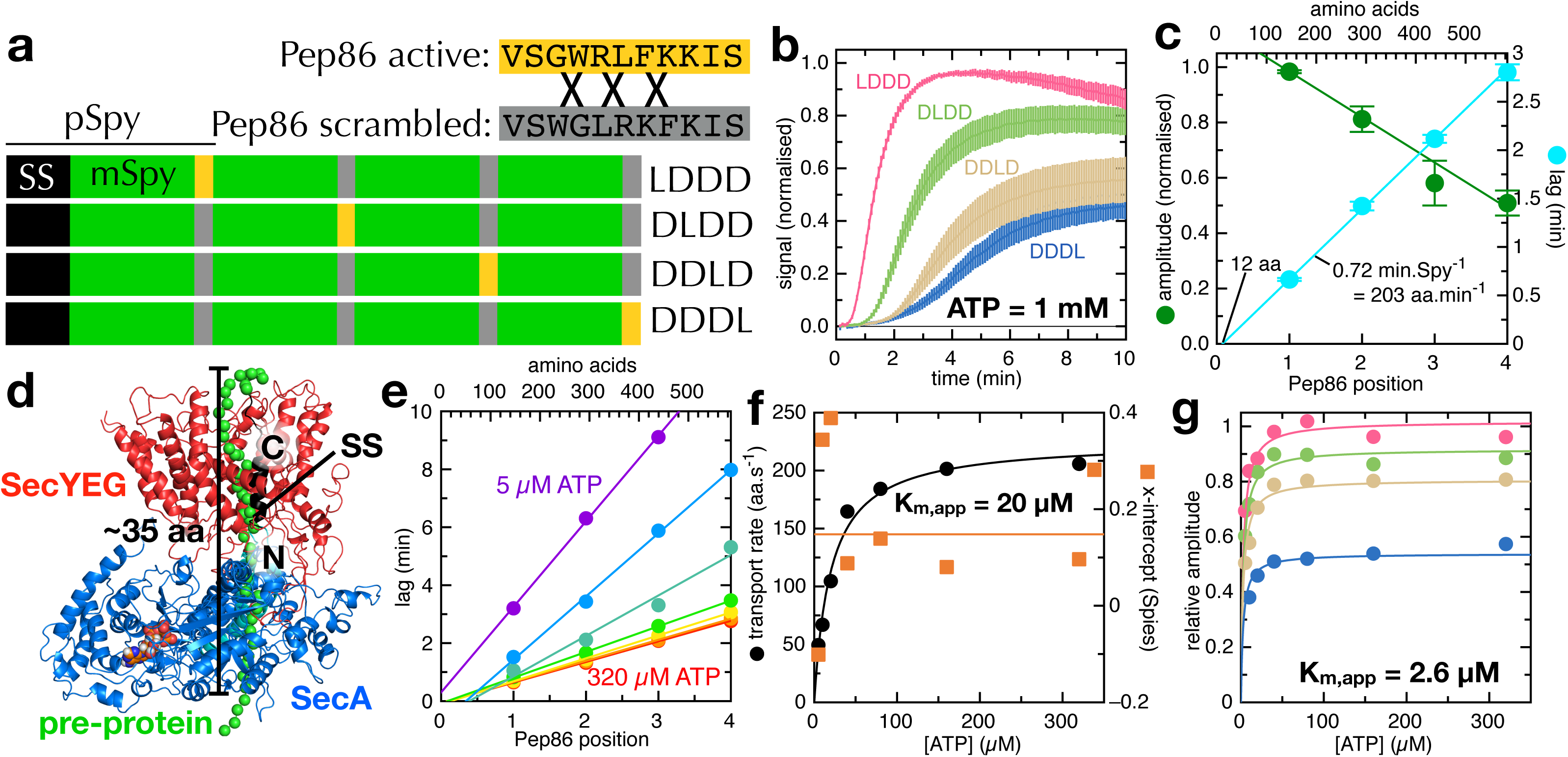
The tandem pSpy-Pep86 series. **a)** Schematic of the tandem pSpy-Pep86 series. **b)** Average transport of 1-2 μM of the pSpy_4x_ series into vesicles containing 10-20 μM 11S, normalised to LDDD for each run. LDDD is pink, DLDD green, DDLD beige and DDDL blue. Error bars are the SEM of 8 repeats. **c)** Normalised amplitude (green circles) and lag (cyan circles) as a function of active Pep86 position (and equivalent in amino acids) for the Spy_4x_ series, extracted from the data in panel **b**. Fits are to straight lines. **d)** Model of SecYEG (red) and SecA (blue) with a pre-protein in the channel (SS in black, mature in green; (Collinson et al., 2015)). Initiation brings ∼35 amino acids of mature domain into the SecY-SecA complex. **e)** Lag as a function of active Pep86 position in the Spy_4x_ series, at a range of ATP concentrations: red = 320 μM, orange = 160 μM, yellow = 80 μM, green = 40 μM, teal = 20 μM, blue = 10 μM and purple = 5 μM. Fits are to straight lines. **f)** Transport rate (black circles) and x-intercept (orange squares) extracted from the fits in panel **d** as a function of ATP concentration. Transport is fitted to the Michaelis-Menten equation (black line), while the orange line is the mean x-intercept. **g)** Amplitude as a function of [ATP] for the Spy_4x_ series. Lines are global fits to a weak binding equation.

From Eq. 1, a plot of lag as a function of n should fit to a straight line with a slope equal to the rate of transport, and y-axis intercept equal to *k*_init_^−1^. The experimental data do indeed fit well to a straight line (Fig. 2c, cyan line), with a slope of 0.72 min.Spy^−1^. Because this value corresponds to n/*k*_step_ for a single pSpy we cannot at this stage distinguish between many fast steps or few slow ones; but as the mature domain of Spy (mSpy; with SS cleaved) is 146 amino acids long it does allow us to determine an average transport rate: 203 amino acids per minute.

The line of best fit in Fig. 2c goes almost straight through the origin, which could be taken to mean that the value of 1/*k*_init_ is very small – i.e. initiation is very fast compared to *k*_step_. However this does not seem consistent with previous data, which did show a slow initiation step (Fessl et al., 2018). Our alternative interpretation is that initiation is accompanied by transport of a short stretch of polypeptide, equivalent to the amount transported by *k*_step_. Indeed, structural evidence suggests that insertion of SS into the lateral gate with its N-terminus facing the cytosol – a key part of initiation (Berg et al., 2004) – brings about 35 amino acids of the mature domain into the SecY-SecA channel (Fig 2d). Therefore, the simplest explanation for these results is that *k*_init_ and *k*_step_ (in the absence of PMF) are effectively the same process. This is consistent with the notion that the catalytic cycle of SecA is primarily regulating the opening and closing of the channel through SecY (Allen et al., 2016): the same widening event permits insertion of the SS during initiation, and diffusion of the pre-protein during transport.

An unexpected observation from the pSpy_4x_ series is that signal amplitude also reduces as the position of the active Pep86 moves later (Fig. 2b and 2c, green circles). This suggests that a significant proportion of *in vitro* transport events initiate and begin transport, but do not reach completion. The most obvious interpretation of this is that it reflects the processivity of transport; indeed, it has previously been reported that SecA dissociates from a different trapped model substrate with an estimated rate constant of 1.2 min^−1^ (Bauer et al., 2014). But as there is nothing to stop reinitiation and additional rounds of transport where the first one fails, this can only explain a slowing of transport with substrate length, not a reduction in final amplitude. To confirm that the slowdown is not somehow an artefact of 11S binding, we also made PLs containing a range of different 11S concentrations and showed that transport of the pSpy_4x_ series is unaffected (Fig. S3f-g).

### The ATP dependence of pre-protein transport

Both initiation and transport of pre-proteins are driven by cycles of ATP binding and hydrolysis in SecA (Economou et al., 1995). The ATP turnover reaction itself has been well characterised in the past (Gold et al., 2007; Robson et al., 2009), but how it is coupled to transport is less well understood. We therefore measured import of each of the Spy_4x_ series at a range of ATP concentrations. All three parameters (lag, λ and amplitude) are affected by ATP concentration, in a similar manner for all four substrates (Fig. 2e-g and S3h). As expected from Eq. 1, the lag remains proportional to the number of Spys before the active Pep86 (and thus the number of steps n) as [ATP] (and thus *k*_step_) is lowered, with the slope of the line becoming steeper (Fig. 2e). The corresponding transport rates and x-intercepts for these data are plotted in Fig. 2f. Rate as a function of [ATP] fits well to the Michaelis-Menten equation, giving an apparent K_m_ for ATP of 20 μM. And consistent with *k*_init_ simply being *k*_step_, the x- or y-intercept is close to the origin and does not change significantly with [ATP], despite *k*_init_ also requiring ATP turnover. Note that as mentioned above, determining lag accurately becomes more difficult for low values of λ, hence the scatter at low [ATP].

Both amplitude and λ vs [ATP] also fit well to the Michaelis-Menten equation for all four substrates (Fig. 2g and Fig. S3h). The K_m_ for ATP determined from λ (15.4 μM, obtained by globally fitting all four data sets) is very similar to that determined from lag, and fairly close to the value of 46 μM determined for the ATPase activity of translocating SecA (Robson et al., 2009). The small discrepancy perhaps reflects the fact that K_m_ determined here only reports on successfully translocated pre-proteins, whereas bulk ATPase activity includes all SecA. For amplitude (Fig. 2g), the apparent K_m_ for ATP is much lower, at 2.6 μM; the discrepancy presumably arising from differences in the way the two parameters respond to ATP concentration (see below).

### Evaluating possible transport models numerically using Berkeley Madonna

The ‘single exponential plus lag’ equation used thus far (Fig. 1a) is straightforward to fit and describes each data set with reasonable accuracy, however it is not immediately evident what λ actually corresponds to in physical terms. Furthermore, the fits deviate significantly from the data at the point where the lag and exponential meet (which we will refer to as the ‘start phase’) – the part of the curve that should contain information about how the motor is distributed along its substrate (McClelland et al., 2005). We therefore sought to fit the data more directly to physical models of transport using numerical integration techniques.

In a biochemical reaction scheme such as the one in Fig. 1b, the concentration of each component changes as a function of time, dependent on the processes that populate it and depopulate it. Because these processes are themselves concentration dependent, the overall reaction can be described by a set of differential equations. Analytical solutions to such problems become highly complex even for fairly simple reaction schemes, so to evaluate different transport models we used numerical integration, as implemented by the software package Berkeley Madonna. In this method, the complete set of differential equations for a given model is defined (see section S2), along with all rates, and the initial concentrations of each species. Next, each concentration is recalculated in very small time increments, and formation of the measured component – in this case NanoLuc – determined as a function of time. These simulated data can then be compared to experimental data, varying the unknown values to try to obtain a reasonable fit.

The model in Fig. 1b is defined for Berkeley Madonna in section S2.1. For simplicity, two additional assumptions are made: that step (*i*) is at equilibrium when the reaction starts, and that NanoLuc formation is instant. The first is reasonable, given that the assay setup includes an 8 min incubation step prior to the addition of ATP and *k*_on_ is of the order of 1.25 μM^−1^.min^−1^ (see above), while the latter is effectively true under the conditions used here (see Fig. 1e). The value *k*_init_ is also set to equal *k*_step_, as per the results above, however the same results are produced if *k*_init_ is set very fast and n is increased by 1.

Figure 3 shows the concentration of each translocation intermediate as a function of time when this model is simulated for n = 5. Initiated complex (red line) begins to form immediately, then eventually decays to a steady state as the pre-formed SecYEG-SecA-pre-protein complex is depleted. The next intermediate (orange line; where the first ∼1/5 of mSpy is inside the vesicle) begins to form after a short delay, as it must wait for initiation. Formation of each subsequent translocation intermediate, with progressively more of the pre-protein inside the PL (yellow (2/5), green (3/5) and teal (4/5) in Fig. 3a), is further offset – as expected for an n-step sequential model. The final product (imported pSpy-Pep86; blue line in Fig. 3a) starts to accumulate after the characteristic lag.

**Figure 3:**
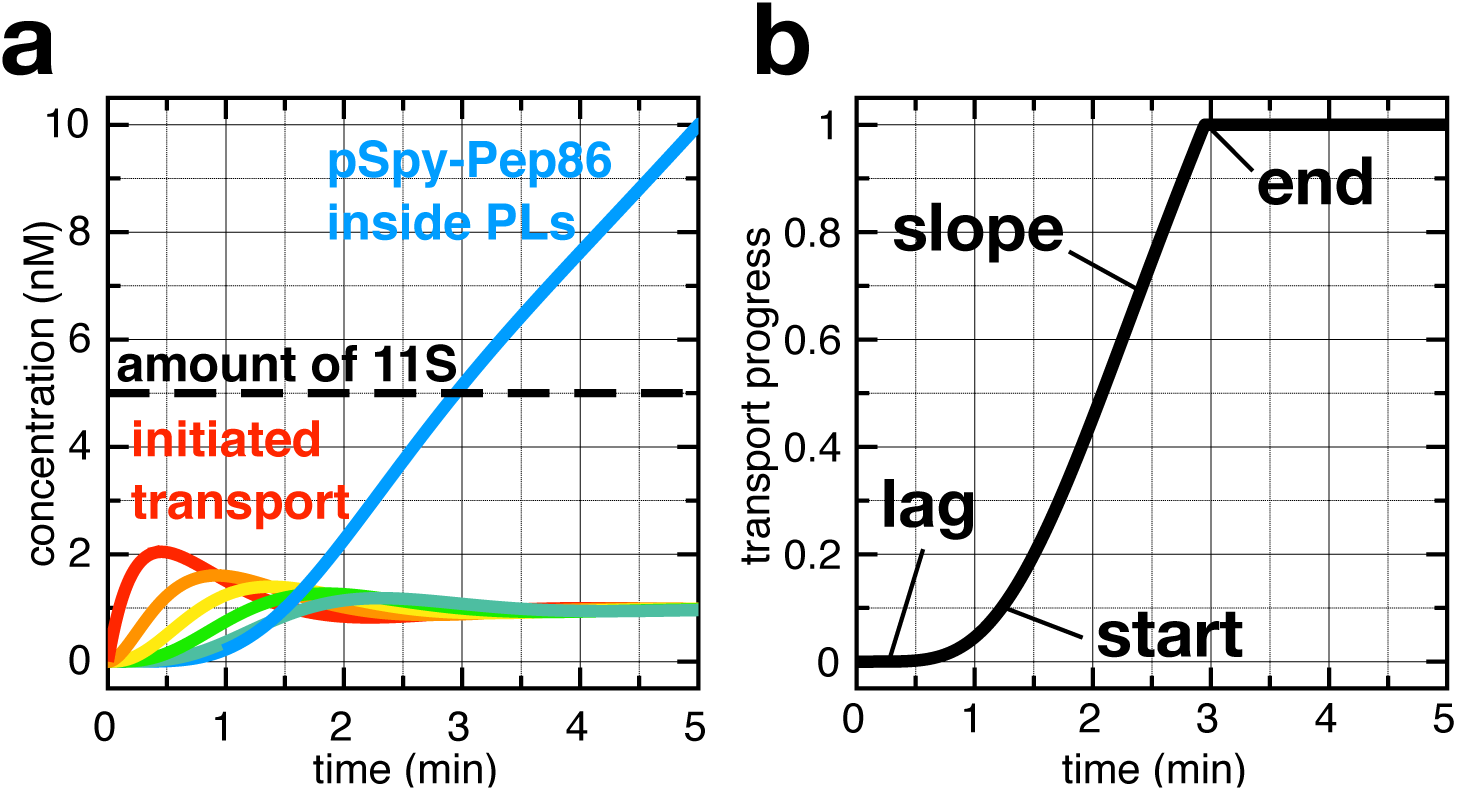
Developing the model of NanoLuc transport. **a)** Simulated concentration of each transport intermediate (for n = 5) as a function of time for the model in Fig. 1b. Red is the initiated transport reaction; orange, yellow, green and teal are progressively later intermediates; and blue is internalised pSpy-Pep86. The total concentration of 11S corresponding to 40 μM internal concentration is shown as a dotted line. Other parameters are: *k*_step_ = 2.5, [pSpy-Pep86] = 1 μM, [SecYEG] = 8.4 nM, [11S] = 5 nM, *k*_on_ = 1.25 and *k*_off_ = 0.75. **b)** Simulated NanoLuc signal from the reaction in panel **a**.

Because of the intrinsic design requirements of the NanoLuc assay, the total amount of 11S present (∼1-5 nM) is a tiny fraction of the total amount of pSpy-Pep86 (generally 1-2 μM) – as illustrated with a black dotted line in Fig. 3a. Therefore, when the luminescent signal produced by the assay is modelled (Fig. 3b) it mirrors the internal pSpy-Pep86 concentration until all the 11S has been saturated, at which point it cannot increase any further. This gives all simulated transport curves the same basic shape: a lag phase characteristic of multi-step reactions; a start phase, where NanoLuc begins to be formed; a straight line slope phase, achieved once all the intermediates reached a steady state; followed by an abrupt end, as 11S runs out (see Fig. 3b).

### Refining the transport model to fit the experimental data

It is fairly clear from Fig. 3b that the initial model (Fig. 1b) is insufficient to describe the experimental data – and not just because of the shape of the end phase. The modelled transport reaction always ends at exactly the same point – when 11S runs out – regardless of what values are chosen for the other parameters. Yet, we know this is not the case: slowing *k*_step_ (by lowering [ATP]; Fig. 2e-f), increasing n (with the tandem pSpy-Pep86 series; Fig. 2b-c) or even lowering [pSpy-Pep86] (Fig. 1d) can all reduce the maximum signal amplitude despite [11S] remaining constant.

As neither pSpy-Pep86 nor 11S are running out, there must be another reason for this behaviour. The most plausible alternative explanations for the reaction to end are either the PLs are becoming ‘full’ as the reaction progresses – i.e. enough pSpy-Pep86 has accumulated inside the PLs that SecA is no longer able to push against its concentration gradient – or the SecYEG import sites are ceasing to function over time, due to translocation stalling, or release failure – preventing multiple pre-protein turnovers.

While we cannot rule out some depletion of import sites over time, modelling these effects and systematically altering the parameters quickly reveals that SecYEG depletion cannot account the observed tailoff in signal (Figs. S4-5). It therefore seems inevitable that transport is slowing due to the building pSpy-Pep86 concentration gradient across the membrane. This would also explain why amplitude shows such a strong dependence on pSpy-Pep86 concentration (Fig. 1d), despite substrate ostensibly not limiting the reaction.

This kind of dependence on the pre-protein gradient across the membrane is inconsistent with a tightly coupled motor, in which one ATP turnover transports a fixed stretch of amino acids, but is exactly what one might expect for a Brownian ratchet. This can be illustrated using our previously proposed Brownian ratchet model (Ahdash et al., 2019; Allen et al., 2016), summarised in Fig. 4a. Each ATP turnover has a probability of leading to transport (p_res_) based on the relative rates of outward and inward diffusion. Because the diffusion rates are dependent on the concentrations of pre-protein on the two sides of the membrane, the overall rate of transport should be too. Effectively, p_res_ decreases as the concentration gradient builds until it reaches 0 and the transport reaction ends.

**Figure 4:**
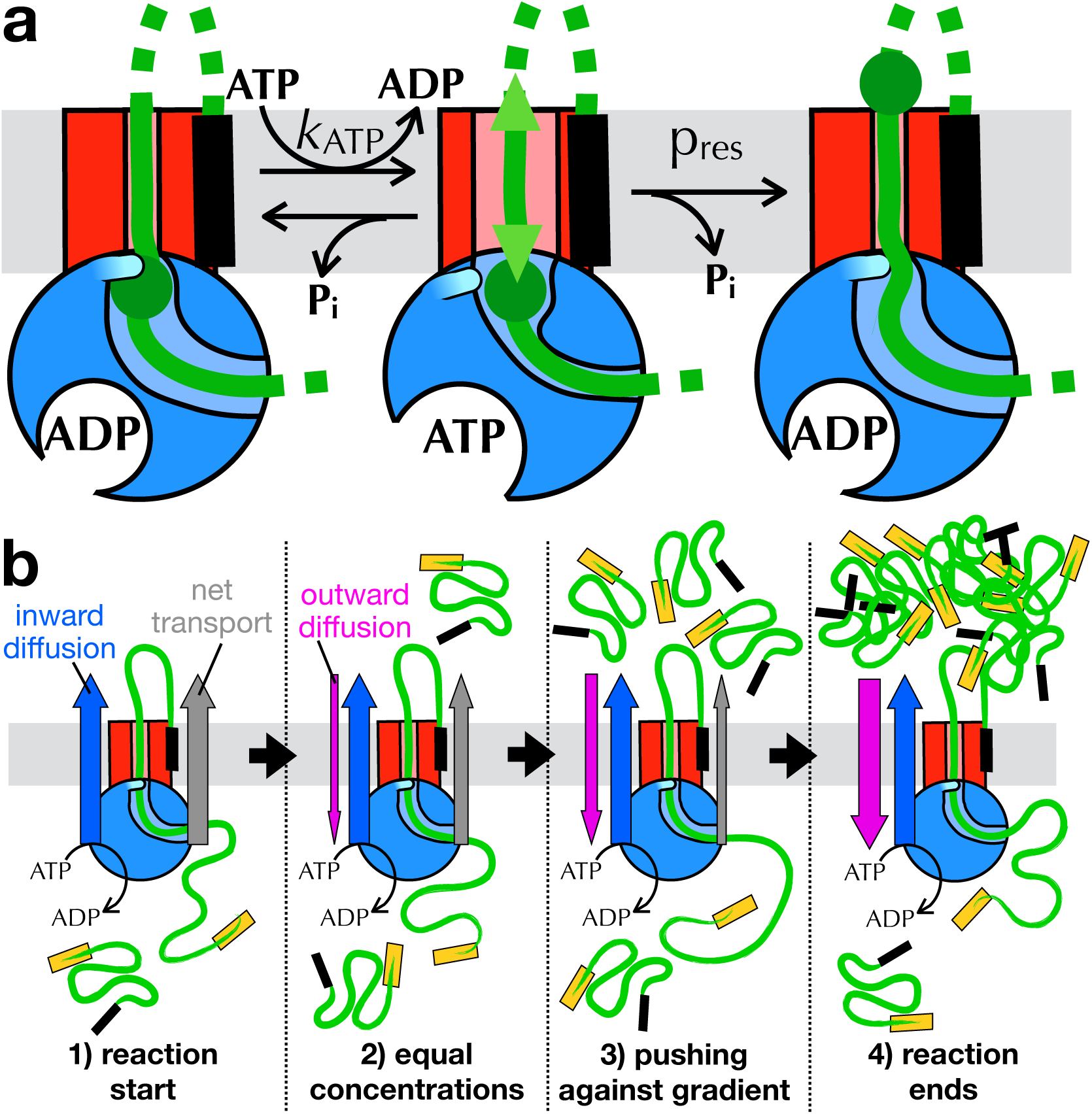
Modelling a Brownian ratchet. **a)** Schematic of the action of ATP in the Brownian ratchet model of pre-protein transport, with components coloured as in Fig. 1b. In the ATP bound state (middle), blockages at the entrance to SecY have a probability of resolving (p_res_) determined by the corresponding diffusion through the channel. **b)** Schematic illustrating the effect of concentration gradient on transport, with components coloured as in Fig. 1b. The net rate of transport (grey arrow) is the sum of the rates of forward (blue arrow) and reverse (pink arrow) diffusion.

To model this effect numerically, it helps to consider the relative diffusion rates of the pre-protein as it is being transported. These are illustrated in Fig. 4b: at the beginning of the transport reaction (**1**) there is no pre-protein inside the PL, so there is only forward diffusion into the PL lumen (blue arrow). This therefore acts as a driver of net transport (grey arrow). As pre-protein accumulates inside the PLs, backward diffusion (pink arrow) becomes a factor, however this is strongly inhibited by the SecA ratchet; thus, when the external and internal concentrations are equal (**2**) transport still proceeds at close to its initial rate. Eventually, however, internal pre-protein accumulates (**3**) to the point where the outward and inward diffusion rates are equal (**4**), and transport stops. This equilibrium is determined by how strongly SecA inhibits backsliding. The fact that concentration gradients influence transport rates is a fundamental feature of thermodynamics, but not one that usually manifests in molecular motors that run along polymers, other perhaps than those that package DNA into viruses.

### What does the amplitude mean?

When the above effect is incorporated into the Berkeley-Madonna model – by scaling *k*_step_ linearly with the ratio of the internal and external concentrations of pre-protein (S2.4 and S2.5) – it fixes many of the discrepancies between the simulated and experimental data (Fig. S6; see also below). But, it still does not explain why signal amplitude goes down as the concentration of ATP is lowered (simulated by lowering *k*_step_; Fig. S6c). To understand this, it helps to consider what the amplitude means physically. As described above, the transport reaction slows as the biased rates of polypeptide diffusion approach parity (Fig. 4b). Seen this way, amplitude is a measure of how great a concentration imbalance SecA can sustain – i.e. SecA motor strength. Lowering [ATP] compromises the coupling between the hydrolytic cycle of ATP and channel opening (Fig. 4a), weakening the motor. As motor speed and ‘strength’ are two subtly different parameters, it makes sense that they saturate at different ATP concentrations (see above; Fig. 2f-g).

An absolute value for SecA strength can be estimated directly from the plot of transport signal amplitude as a function of pSpy-Pep86 concentration (Fig. 1d, replotted in Fig. S7): amplitude increases linearly with pSpy-Pep86 concentration up to the point where 11S becomes limiting, when importing more pSpy-Pep86 has no further effect; at this point the interior concentration of pSpy-Pep86 must be at least that of 11S. As these reactions were performed with PLs containing 40 μM 11S and saturate at 0.4 ± 0.2 μM external pSpy concentration (Fig. S7; error is the SEM from four repeats), we estimate that SecA with saturating ATP can push against a pSpy concentration gradient of the order of 100x. This value will presumably vary between different preproteins, and be affected by other external factors that promote or inhibit pre-protein transport *in vivo*, such as the PMF.

### Using the Berkeley Madonna model to estimate elementary step size

The model incorporating slowdown is the simplest one capable of describing all the experimental data – but it does not mean no other factors contribute. Two in particular are worth mentioning: firstly the fact that the PLs are not all the same size will change the shape of the end phase (see Fig. S8), meaning we cannot determine an accurate value for slowdown. And secondly static disorder – the phenomenon whereby different, ostensibly identical motors can transport at different rates (Bianco et al., 2001; Park et al., 2010) – can lead to an underestimation of the true number of steps (Fig. S9). For example, ensemble measurements on the helicase PcrA overestimate step size about 4-fold compared to the true value determined by single molecule (Park et al., 2010).

Notwithstanding these effects, the shape of the beginning of the transport curve should allow us to estimate of the number of steps that make up transport. To do this we fitted the pSpy-Pep86 data manually for a range of different values of n, adjusting *k*_step_ and its concentration dependence to best fit the start phase. The results of this for n = 2 to n = 9 are shown in Fig. 5. Overall, the best fit to the start phase gives a value of 5 steps per mSpy (excluding *k*_init_), with a *k*_step_ of about 2.8 min^−1^. This equates to about one step every ∼30 amino acids (Fig. 5). Although slightly better fits can be obtained if we allow all the other parameters to vary as well, the start phase is always too shallow for n < 2, and too sharp for n > 8. Therefore, we conclude that transport is best described by a relatively small number of steps.

**Figure 5:**
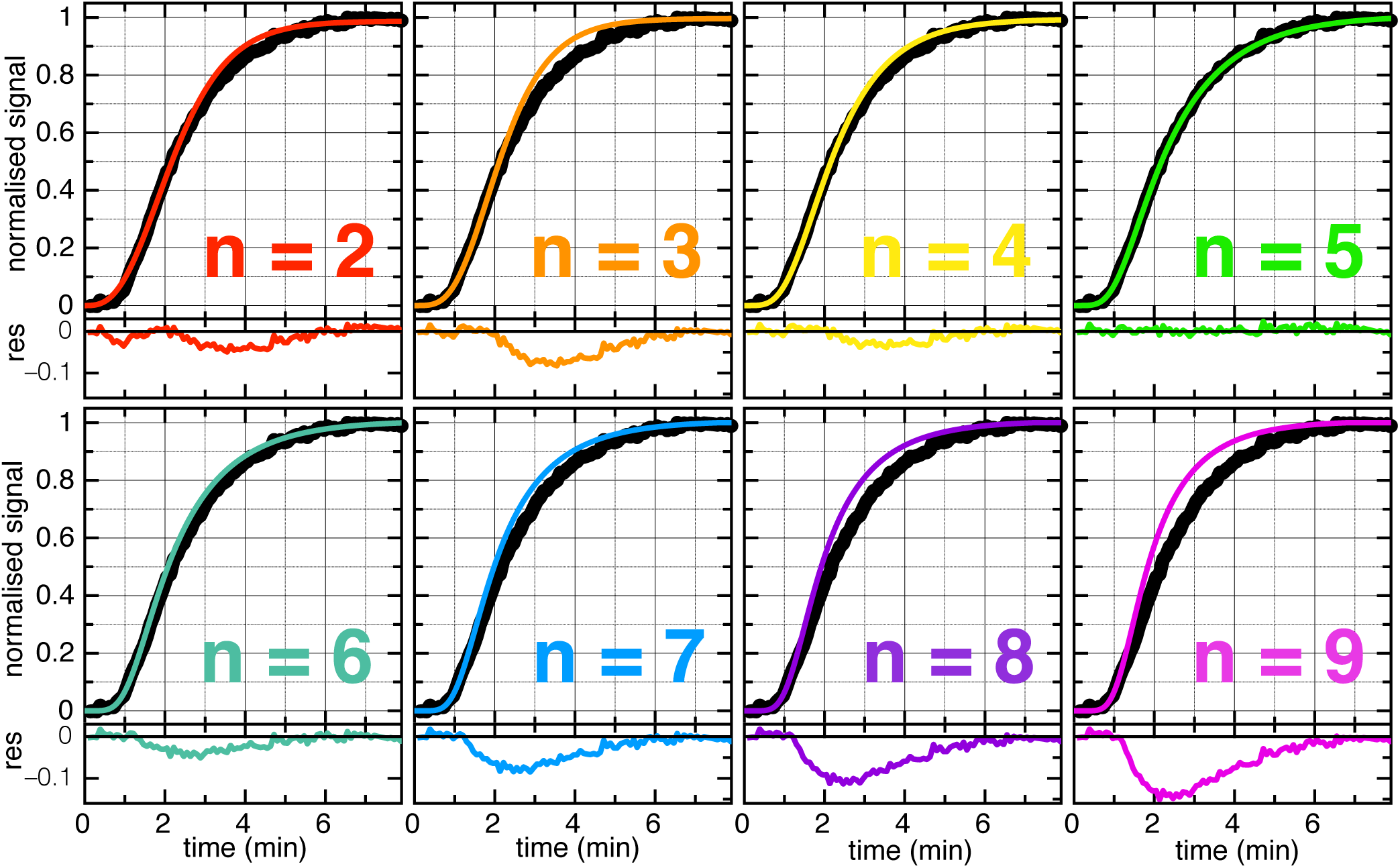
The final transport model. Transport of pSpy-Pep86 (black dots; average of 3 repeats) with best fits to: n = 2 (red; *k*_step_ = 0.4 min^−1^, slowdown = 2433), n = 3 (orange; *k*_step_ = 0.86 min^−1^, slowdown = 1249), n = 4 (yellow; *k*_step_ = 1.84 min^−1^, slowdown = 345), n = 5 (green; *k*_step_ = 2.83 min^−1^, slowdown = 196), n = 6 (teal; *k*_step_ = 3.5 min^−1^, slowdown = 196), n = 7 (blue; *k*_step_ = 4.2 min^−1^, slowdown = 196), n = 8 (purple; *k*_step_ = 4.9 min^−1^, slowdown = 196) and n = 9 (magenta; *k*_step_ = 5.7 min^−1^, slowdown = 196). Other parameters are: [SecYEG] = 8.4 nM, [11S] = 5 nM, *k*_on_ = 1.25 min^−1^ and *k*_off_ = 0.75 min^−1^. Residuals are shown below each graph.

The measured *k*_cat_ of the SecA ATPase activity when transporting pSpy is ∼850 min^−1^ for *in vitro* transport with purified components (Fig. S10); not too far from the previously determined value of 450 min^−1^ for proOmpA (Robson et al., 2009). This corresponds to around 300 ATP turnovers per step. In the Brownian ratchet transport model (Fig. 4a) this would mean p_res_ – i.e. the probability of any given ATP turnover producing a transport event – is only 0.33% even before slowdown from the concentration gradient is accounted for. And even if the Sec system were to display static disorder similar to PcrA, leading e.g. to a five-fold overestimate of step size, this would still be equivalent to steps of 6 amino acids with an average p_res_ of 1.67%. For a classical power stroke mechanism, meanwhile, we would expect this value to approach 100%. Of course, given the variability of pre-protein sequences there is no particular reason to believe that all steps are equally sized, or that they traverse the channel at exactly the same rate – these numbers therefore represent averages, and will doubtless vary between different pre-proteins as well.

## Discussion

The advent of an assay capable of measuring protein transport accurately, with a time resolution of seconds, has opened the door to investigating the process on a detailed functional as well as a structural level. Using the NanoLuc assay, we have here built up a thorough understanding of how each transport parameter affects the measured transport signal and developed two ways to analyse them. Fitting to a simple model requires no specialist software and gives two useful parameters: lag, which corresponds to the minimum time required for transport; and amplitude, which corresponds to the strength of the SecA motor – provided the pre-protein concentration is not enough to saturate all the 11S. A complete numerical integration solution, meanwhile, validates the overall model and provides estimates for each individual rate constant. It should be noted that lag is relatively independent of experimental variables, and is thus a robust measure of transport rate. Amplitude, meanwhile, is highly susceptible to experimental error (see Fig. S7); care must therefore be taken when using it to interpret differences in motor strength.

Our results strongly support the notion that ATP hydrolysis is indirectly, not directly coupled to protein transport – as in a Brownian ratchet rather than a power stroke. In particular, the strong dependence of transport rate on the cross-membrane pre-protein concentration can only be explained if diffusion is an integral part of the mechanism. Initiation, although clearly a critical part of the mechanism for recognising genuine Sec substrates, does not seem to contribute appreciably to the overall kinetics of transport. Indeed, if anything the first few amino acids of mSpy are transported faster than the rest (Fig. 2c,e). A likely explanation for this is that insertion of the signal sequence into the LG of SecA – oriented to keep the positively charged N-terminus in the cytosol and bring the narrow, hydrophilic C-terminus through the channel (Fig. 2d) – provides an extra driving force to pull the first amino acids across the membrane.

The net transport rate of ∼200 amino acids per minute determined here is similar to one previous estimate of transport rate, for fluorescently labelled proOmpA into IMVs (Tomkiewicz et al., 2006), but roughly 10-fold slower than translocation of unlabelled proOmpA into PLs (Fessl et al., 2018). This can partly be ascribed to the nature of the substrate: proOmpA was originally chosen as a model translocation substrate precisely because it is secreted very efficiently. However it should also be noted that the NanoLuc assay reports only on successful transport events, whereas the single molecule assay in Fessl *et al.* uses movement of the plug domain of SecY as a proxy for transport. Therefore, the higher rates reported in Fessl *et al*. may also partially reflect a fraction of initiated, but subsequently aborted events.

Thorough studies of the kinetics of protein import have been performed previously, and are usually fitted either qualitatively (Tomkiewicz et al., 2006) or to a single exponential (Bariya and Randall, 2018). Our conclusions here allow these results to be interpreted in a new light. Such assays are almost always performed with a total reaction volume far greater than the internal lumen volume of the PLs or IMVs into which it is being transported. So just as here, it seems likely that the transport is reaching completion when the concentration gradient across the membrane equals the force provided by SecA. The exact amount of pre-protein this corresponds to will therefore depend on the strength of the SecA motor, but also the total internal volume of the reaction vesicles. The rate at which it approaches this, meanwhile, is effectively the rate of transport divided by the amount inside at equilibrium. As these are to some extent correlated – good Sec substrates transport fast, but more can enter the PLs – this value might be expected to be relatively insensitive to reaction conditions.

It is interesting to note that the slowing down of the transport reaction as the vesicles become fuller is in effect an artefact of measuring transport into PLs with a limited internal volume. In the context of a living cell, secreted pre-proteins do not accumulate statically; rather the signal peptide is cleaved off and they are either folded or transported away. However, this measurement artefact is common to all previous pre-protein transport assays that measure transport into vesicles – i.e. nearly all of them – and would not have been spotted without the high resolution of the NanoLuc assay and detailed dissection of its mechanism.

The slowing effect from vesicle fullness goes perhaps some way towards explaining why transport assays performed *in vitro* are so much slower than the rates expected *in vivo* (Collinson et al., 2015; Cranford-Smith and Huber, 2018). But it cannot explain a predicted difference of nearly 2 orders of magnitude (Collinson et al., 2015; Cranford-Smith and Huber, 2018). A clue to this may come from the extremely low probability that any given ATP turnover event gives rise to transport *in vitro* (p_res_ = 0.33%; see Fig. 4a). This value seems implausibly low, so it is very likely that other factors will increase this value substantially *in vivo*. These might include auxiliary drivers of transport, particularly the PMF (Brundage et al., 1990; Schiebel et al., 1991), and some of the many other proteins that associate with the Sec system such as SecDF, PpiD and YfgM (Jauss et al., 2019; Pogliano and Beckwith, 1994). It does, however, makes sense that all these factors affect p_res_, not ATP turnover itself: it is far easier to add additional driving forces to ratchet than to a directly coupled motor.

One additional factor we believe will prove particularly critical to understanding the slow *in vitro* transport rates is the folding state of the pre-protein, which is known to be important for enabling transport (Corey et al., 2019; Tsirigotaki et al., 2018). Chaperones generally capture pre-proteins *in vivo* as they are translated and deliver them to the membrane in an optimally translocation competent state. *In vitro*, meanwhile, pre-proteins are diluted out of urea, and so have far more opportunity to form folding intermediates that delay transport. Without extra assistance, the diffusion-based transport motor has little power to unfold pre-proteins: instead it must wait for a sponteneous unfolding event then trap them in an unfolded state within the channel. It seems plausible that this effect could slow transport by at least an order of magnitude.

The twin developments described here: an assay that generates high quality transport data and a fitting process capable of describing it, together provide the first fully quantitative framework for understanding the mechanism of ATP-driven transport through Sec. We anticipate that the experimental and data analysis approaches developed here will be very useful in the future both for furthering our understanding of the bacterial Sec machinery, but also far more broadly to study many other membrane transport processes.

## Methods

### Reagents

All reagents for transport assays, including SecYEG, SecA, pSpy and PLs were produced exactly as described previously (Pereira et al., 2019).

The Spy_4x_ series was constructed by first ordering 3 fragments of Spy containing 2 tandem repeats of the mature region, each with a different combination of ‘light’ (L, active Pep86, VSGWRLFKKIS) or ‘dark’ (D, inactive pep86, VSWGLRKFKIS) Pep86 sequences in the following combinations: DD, LD and DL, where for example, DD contained two inactive HiBiT sequences at the C terminus of each mature region of Spy (produced by the GeneArt Gene Synthesis service, Thermo Fisher Scientific). For cloning purposes, each fragment began with residue A24 of pSpy and ended with a GSG linker immediately following the second Pep86 sequence (sequence 1 in Fig. S2). The fragment was cloned into pBAD-pSpy-V5-pep86-TEV-His ((Pereira et al., 2019); sequence 2 in Fig. S2) using site directed ligase independent mutagenesis (Chiu et al., 2004). More specifically, the fragments were cloned in the place of Spy-V5, using the same primers to introduce a ZraI site (GACGTC) immediately after the fragment and before the TEV cleavage site of the template, to give pBAD-(LL, LD or DL)-ZraI-TEV-His (sequence 3 in Fig. S2). The synthesised fragments were then amplified using linear PCR and cloned into the ZraI site of pSpy LL, LD and DL to give DDDD, LDDD, DLDD, DDLD and DDDL (Sequence 4 in Fig. S2).

### NanoLuc formation assays

NanoLuc formation was measured at 25 °C in a BioTek Synergy Neo2 plate reader. A dilution series of pSpy-Pep86 was prepared in the wells using TKM with furimazine (to 1/500) and Prionex (to 0.125%), with a volume of 100 μl per well. Reactions were started by injecting 25 μl 11S at 1 nM in TKM (to give 200 pM final), then shaken for 2 s and the luminescence monitored with no emission filter.

### NanoLuc transport experiments

Transport experiments were performed exactly as in (Pereira et al., 2019).

### Data analysis

Initial data processing was performed using pro Fit 7 (Quansoft). Raw data before the addition of ATP were fitted to the single molecule plus lag model (Section S1.1), and the fits subtracted to give transport signal (see Fig. S1a). Corrected data were then fitted to the same model, to give lag, λ and amplitude for transport. Models for Berkeley Madonna are listed and described in detail in section S2.

### Dynamic light scattering

DLS measurements were performed in a Malvern zetasizer nano, using the same PLs as for transport (extruded to 400 nm). Vesicle population as a function of size were determined using the built in instrument software (Zetasizer).

### ATPase assays

ATPase assays wer performed exactly as in (Gold et al., 2007)

## Notes

### Competing Interest Statement

The authors have declared no competing interest.

