## Supplementary text for "Refined measurement of SecA-driven protein transport reveals indirect coupling to ATP turnover"

**Supplementary Information**

**Section S1. pro Fit models**

**S1.1. Single exponential plus lag**

```
function Exp_plus_lag(amp, k, C, lag: real);

inputs
amp := 0, active;
k := 0, active;
C := 0, inactive;
lag := 0, active;

var
xapp: real;

begin
xapp := x - lag;

if xapp < 0
	then y := C

else
y := amp * (1 - exp(-xapp * k)) + C;

end;
```

**S1.2. Weak binding equation**

function WeakBinding(Kd, F0, Fb: real);

inputs

Kd := 2, active;

F0 := 0, active;

Fb := 1, active;

var

PA, PB: real;

begin

PB := x/(x + Kd);

PA := 1 - PB;

y := (F0 * PA) + (Fb * PB);

end;

**S1.3. Michaelis Menten equation**

function MichaelisMenten(Km, kcat: real);

inputs

Km := 2, active;

kcat := 10, active;

begin

y := (kcat * x) / (Km + x);

end;

**Section S2. Berkeley Madonna Models**

This section contains all the Berkeley Madonna models used to fit the data. For each model beyond the first, changes from the template model are highlighted in bold.

**S2.1 – Basic model**

Simulating the model in Fig. 1b (highlighted pink). Parameters are as follows: A is the Sec machinery; B is pre-protein; C is pre-initiation complex; D[0] is post-initiation complex; D[1] to D[n-1] are translocation intermediates; D[n] is translocated pre-protein; and Lg is 11S.

METHOD RK4

STARTTIME = 0

STOPTIME = 10

DT = 0.001

{*** Initial parameters ***}

{rate constants (in µM and min, as appropriate)}

k_on = 1.25

k_off = 0.75

k_step = 5

k_init = k_step

{supplied concentrations (µM)}

Lg = 0.005 {11S concentration (final)}

A0 = 0.0084 {starting SecYEG concentration}

B0 = 1 {starting pre-protein concentration}

{additional parameters}

n = 5 {number of time k_step is repeated}

brightness = 1/Lg {how much signal is produced per nLuc}

{*** Reaction setup ***}

{initiate concentrations}

b_quad = A0 + B0 + (k_off / k_on)

INIT C = (b_quad - SQRT((b_quad * b_quad) - (4 * A0 * B0))) / 2

INIT A = A0 - C

INIT B = B0 - C

INIT D[0..n] = 0

{*** Differential equations ***}

d/dt (A) = (C * k_off) - (A * B * k_on) + (D[n-1] * k_step)

d/dt (B) = (C * k_off) - (A * B * k_on)

d/dt (C) = (A * B * k_on) - (C * k_off) - (C * k_init)

d/dt (D[0]) = (C * k_init) - (D[0] * k_step)

d/dt (D[1..n-1]) = (D[i-1] * k_step) - (D[i] * k_step)

d/dt (D[n]) = (D[n-1] * k_step)

{*** Output ***}

signal = min(D[n], Lg) * brightness

**S2.2 – SecYEG unable to perform multiple turnovers**

Same as the basic model, but SecYEG (A) no longer becomes available once transport of one pre-protein is complete. The missing parameter is shown struck through; note that for this model to function in Berkeley Madonna it must be deleted.

METHOD RK4

STARTTIME = 0

STOPTIME = 10

DT = 0.001

{*** Initial parameters ***}

{rate constants (in µM and min, as appropriate)}

k_on = 1.25

k_off = 0.75

k_step = 5

k_init = k_step

{supplied concentrations (µM)}

Lg = 0.005 {11S concentration (final)}

A0 = 0.0084 {starting SecYEG concentration}

B0 = 1 {starting pre-protein concentration}

{additional parameters}

n = 5 {number of time k_step is repeated}

brightness = 1/Lg {how much signal is produced per nLuc}

{*** Reaction setup ***}

{initiate concentrations}

b_quad = A0 + B0 + (k_off / k_on)

INIT C = (b_quad - SQRT((b_quad * b_quad) - (4 * A0 * B0))) / 2

INIT A = A0 - C

INIT B = B0 - C

INIT D[0..n] = 0

{*** Differential equations ***}

d/dt (A) = (C * k_off) - (A * B * k_on) ~~+ (D[n-1] * k_step)~~

d/dt (B) = (C * k_off) - (A * B * k_on)

d/dt (C) = (A * B * k_on) - (C * k_off) - (C * k_init)

d/dt (D[0]) = (C * k_init) - (D[0] * k_step)

d/dt (D[1..n-1]) = (D[i-1] * k_step) - (D[i] * k_step)

d/dt (D[n]) = (D[n-1] * k_step)

{*** Output ***}

signal = min(D[n], Lg) * brightness

**S2.3 – SecYEG becoming inactive over time**

This model modifies the basic model allowing translocating SecYEGs to decay over time with the rate k_Sec_death.

METHOD RK4

STARTTIME = 0

STOPTIME = 10

DT = 0.001

{*** Initial parameters ***}

{rate constants (in µM and min, as appropriate)}

k_on = 1.25

k_off = 0.75

k_step = 5

k_init = k_step

**k_Sec_death = 1**

{supplied concentrations (µM)}

Lg = 0.005 {11S concentration (final)}

A0 = 0.0084 {starting SecYEG concentration}

B0 = 1 {starting pre-protein concentration}

{additional parameters}

n = 5 {number of time k_step is repeated}

brightness = 1/Lg {how much signal is produced per nLuc}

{*** Reaction setup ***}

{initiate concentrations}

b_quad = A0 + B0 + (k_off / k_on)

INIT C = (b_quad - SQRT((b_quad * b_quad) - (4 * A0 * B0))) / 2

INIT A = A0 - C

INIT B = B0 - C

INIT D[0..n] = 0

{*** Differential equations ***}

d/dt (A) = (C * k_off) - (A * B * k_on) + (D[n-1] * k_step)

d/dt (B) = (C * k_off) - (A * B * k_on)

d/dt (C) = (A * B * k_on) - (C * k_off) - (C * k_init)

d/dt (D[0]) = (C * k_init) - (D[0] * k_step) **- (D[0] * k_Sec_death)**

d/dt (D[1..n-1]) = (D[i-1] * k_step) - (D[i] * k_step) **- (D[i] * k_Sec_death)**

d/dt (D[n]) = (D[n-1] * k_step)

{*** Output ***}

signal = min(D[n], Lg) * brightness

**S2.4 – Transport model with slowing as a pre-protein enters the PLs**

In this model, the basic model (S2.1) is modified to slow transport down (by the factor 'slowdown') as pre-protein is transported into the PLs, to a minimum value of 0**.** Brightness is also uncoupled from [11S], allowing the amplitude to be scaled freely.

METHOD RK4

STARTTIME = 0

STOPTIME = 10

DT = 0.001

{*** Initial parameters ***}

{rate constants (in µM and min, as appropriate)}

k_on = 1.25

k_off = 0.75

k_step**_init** = 5

k_init = k_step_**init**

{supplied concentrations (µM)}

Lg = 0.005 {11S concentration (final)}

A0 = 0.0084 {starting SecYEG concentration}

B0 = 1 {starting pre-protein concentration}

{additional parameters}

n = 5 {number of time k_step is repeated}

brightness = **200** {how much signal is produced per nLuc}

{*** Reaction setup ***}

{initiate concentrations}

b_quad = A0 + B0 + (k_off / k_on)

INIT C = (b_quad - SQRT((b_quad * b_quad) - (4 * A0 * B0))) / 2

INIT A = A0 - C

INIT B = B0 - C

INIT D[0..n] = 0

**{modulating k_step}**

**slowdown = 300**

**k_step = k_step_init * max(0, (1 - (slowdown * D[n] / B)))**

{*** Differential equations ***}

d/dt (A) = (C * k_off) - (A * B * k_on) + (D[n-1] * k_step)

d/dt (B) = (C * k_off) - (A * B * k_on)

d/dt (C) = (A * B * k_on) - (C * k_off) - (C * k_init)

d/dt (D[0]) = (C * k_init) - (D[0] * k_step)

d/dt (D[1..n-1]) = (D[i-1] * k_step) - (D[i] * k_step)

d/dt (D[n]) = (D[n-1] * k_step)

{*** Output ***}

signal = min(D[n], Lg) * brightness

**S2.5 – Slowing transport of tandem pSpy series**

Here, the model with slowing (S2.4) is modified to give a signal after different amounts of pre-protein have been transported, such that the total pre-protein length is 4n. The first step of the first transport event is also removed, as it is accounted for by initiation.

METHOD RK4

STARTTIME = 0

STOPTIME = 10

DT = 0.001

{*** Initial parameters ***}

{rate constants (in µM and min, as appropriate)}

k_on = 1.25

k_off = 0.75

k_step_init = 5

k_init = k_step_init

{supplied concentrations (µM)}

Lg = 0.005 {11S concentration (final)}

A0 = 0.0084 {starting SecYEG concentration}

B0 = 1 {starting pre-protein concentration}

{additional parameters}

n = 5 {number of time k_step is repeated}

brightness = 200 {how much signal is produced per nLuc}

{*** Reaction setup ***}

{initiate concentrations}

b_quad = A0 + B0 + (k_off / k_on)

INIT C = (b_quad - SQRT((b_quad * b_quad) - (4 * A0 * B0))) / 2

INIT A = A0 - C

INIT B = B0 - C

INIT D[**1**..**4***n] = 0

**INIT LDDD = 0**

**INIT DLDD = 0**

**INIT DDLD = 0**

**INIT DDDL = 0**

{modulating k_step}

slowdown = 300

k_step = k_step_init * max(0, (1 - (slowdown * D[**4***n] / B)))

{*** Differential equations ***}

d/dt (A) = (C * k_off) - (A * B * k_on) + (D[**4***n-1] * k_step)

d/dt (B) = (C * k_off) - (A * B * k_on)

d/dt (C) = (A * B * k_on) - (C * k_off) - (C * k_init)

d/dt (D[**1**]) = (C * k_init) - (D[**1**] * k_step)

d/dt (D[**2**..**4***n-1]) = (D[i-1] * k_step) - (D[i] * k_step)

d/dt (D[**4***n]) = (D[**4***n-1] * k_step)

**d/dt (LDDD) = (D[n-1] * k_step)**

**d/dt (DLDD) = (D[2*n-1] * k_step)**

**d/dt (DDLD) = (D[3*n-1] * k_step)**

**d/dt (DDDL) = (D[4*n-1] * k_step)**

{*** Output ***}

**signal_LDDD = min(LDDD, Lg) * brightness**

**signal_DLDD = min(DLDD, Lg) * brightness**

**signal_DDLD = min(DDLD, Lg) * brightness**

**signal_DDDL = min(DDDL, Lg) * brightness**

**S2.6 – Accounting for vesicle size distribution**

To simulate the effect of vesicle size distribution on the model in S2.1 (Fig. 6a), the population of SecYEG is divided up into 18 subpopulations (surf_pop[], based on calculated surface area), and assigned a fraction of the total 11S (vol_pop[], based on volume). These are all treated as completely separate reactions, using arrays for each component. The final signal is the sum of all the individual signals.

METHOD RK4

STARTTIME = 0

STOPTIME = 10

DT = 0.001

{*** Initial parameters ***}

{rate constants (in µM and min, as appropriate)}

k_on = 1.25

k_off = 0.75

k_step_init = 5

k_init = k_step_init

{supplied concentrations (µM)}

Lg = 0.005 {11S concentration (final)}

A0 = 0.0084 {starting SecYEG concentration}

B0 = 1 {starting pre-protein concentration}

{additional parameters}

n = 5 {number of time k_step is repeated}

brightness = 200 {how much signal is produced per nLuc}

**{*** accounting for vesicle size distribution ***}**

**m = 17 {index of last population}**

**surf_pop[0..m] = 0.008441495**

**surf_pop[1] = 0.03821**

**surf_pop[2] = 0.07591**

**surf_pop[3] = 0.09621**

**surf_pop[4] = 0.09763**

**surf_pop[5] = 0.08880**

**surf_pop[6] = 0.07734**

**surf_pop[7] = 0.06755**

**surf_pop[8] = 0.05965**

**surf_pop[9] = 0.05626**

**surf_pop[10] = 0.05562**

**surf_pop[11] = 0.05328**

**surf_pop[12] = 0.05002**

**surf_pop[13] = 0.04792**

**surf_pop[14] = 0.03857**

**surf_pop[15] = 0.03448**

**surf_pop[16] = 0.02312**

**surf_pop[17] = 0.03100**

**vol_pop[0..m] = 0.002223334**

**vol_pop[1] = 0.01165**

**vol_pop[2] = 0.02681**

**vol_pop[3] = 0.03936**

**vol_pop[4] = 0.04625**

**vol_pop[5] = 0.04871**

**vol_pop[6] = 0.04913**

**vol_pop[7] = 0.04971**

**vol_pop[8] = 0.05083**

**vol_pop[9] = 0.05551**

**vol_pop[10] = 0.06356**

**vol_pop[11] = 0.07051**

**vol_pop[12] = 0.07665**

**vol_pop[13] = 0.08505**

**vol_pop[14] = 0.07928**

**vol_pop[15] = 0.08208**

**vol_pop[16] = 0.06374**

**vol_pop[17] = 0.09896**

{*** Reaction setup ***}

{initiate concentrations}

b_quad = A0 + B0 + (k_off / k_on)

**C0** = (b_quad - SQRT((b_quad * b_quad) - (4 * A0 * B0))) / 2

**INIT A[0..m] = (A0 - C0) * surf_pop[i]**

INIT B = B0 - C**0**

**INIT C[0..m] = C0 * surf_pop[i]**

INIT D[0..n**, 0..m**] = 0

{modulating k_step}

slowdown = 300

k_step**[0..m]** = k_step_init * max(0, (1 - (slowdown * D[n**, i**] / B)))

{*** Differential equations ***}

d/dt (A**[0..m]**) = (C**[i]** * k_off) - (A**[i]** * B * k_on) + (D[n-1**, i**] * k_step**[i]**)

d/dt (B) = (**ARRAYSUM(**C**[*]**) * k_off) - (**ARRAYSUM(**A**[*])** * B * k_on)

d/dt (C**[0..m]**) = (A**[i]** * B * k_on) - (C**[i]** * k_off) - (C**[i]** * k_init)

d/dt (D[0**, 0..m**]) = (C**[j]** * k_init) - (D[0**, j**] * k_step**[j]**)

d/dt (D[1..n-1**, 0..m**]) = (D[i-1**, j**] * k_step**[j]**) - (D[i**, j**] * k_step**[j]**)

d/dt (D[n**,0..m**]) = (D[n-1**, i**] * k_step**[j]**)

{*** Output ***}

NLuc**[0..m]** = min(D[n**,i**], (Lg *** vol_pop[i]**))

signal = **ARRAYSUM(**NLuc**[*])** * brightness

**S2.7 – Modelling static disorder**

Here, the transport model with slowdown (S2.4) is modified to simulate the effect of static disorder. Instead of a single value for k_step_init, we simulate 41 different populations of SecYEG, each with a different k_step_init value, populated according to a Gaussian distribution and ranging from (k_step_avrg_init ÷ disorder) to (k_step_avrg_init × disorder). Note that we initially tried to model fewer populations (11), but this produces artefacts under some conditions.

METHOD RK4

STARTTIME = 0

STOPTIME = 10

DT = 0.001

{*** Initial parameters ***}

{rate constants (in µM and min, as appropriate)}

k_on = 1.25

k_off = 0.75

k_step**_avrg**_init = 5

{supplied concentrations (µM)}

Lg = 0.005 {11S concentration (final)}

A0 = 0.0084 {starting SecYEG concentration}

B0 = 1 {starting pre-protein concentration}

{additional parameters}

n = 5 {number of time k_step is repeated}

brightness = 200 {how much signal is produced per nLuc}

**{*** simulating static disorder ***}**

**disorder = 0**

**k_step_init[0..20] = k_step_avrg_init / (1 + (disorder * (20 - i) / 20))**

**k_step_init[21..40] = k_step_avrg_init * (1 + (disorder * (i - 20) / 20))**

**disorder_pop[0..40] = 0.00152899**

**disorder_pop[1] = 0.00216252**

**disorder_pop[39] = 0.00216252**

**disorder_pop[2] = 0.00300465**

**disorder_pop[38] = 0.00300465**

**disorder_pop[3] = 0.00410118**

**disorder_pop[37] = 0.00410118**

**disorder_pop[4] = 0.00549922**

**disorder_pop[36] = 0.00549922**

**disorder_pop[5] = 0.00724392**

**disorder_pop[35] = 0.00724392**

**disorder_pop[6] = 0.00937401**

**disorder_pop[34] = 0.00937401**

**disorder_pop[7] = 0.0119**

**disorder_pop[33] = 0.0119**

**disorder_pop[8] = 0.0149**

**disorder_pop[32] = 0.0149**

**disorder_pop[9] = 0.0183**

**disorder_pop[31] = 0.0183**

**disorder_pop[10] = 0.0220**

**disorder_pop[30] = 0.0220**

**disorder_pop[11] = 0.0261**

**disorder_pop[29] = 0.0261**

**disorder_pop[12] = 0.0303**

**disorder_pop[28] = 0.0303**

**disorder_pop[13] = 0.0346**

**disorder_pop[27] = 0.0346**

**disorder_pop[14] = 0.0389**

**disorder_pop[26] = 0.0389**

**disorder_pop[15] = 0.0429**

**disorder_pop[25] = 0.0429**

**disorder_pop[16] = 0.0464**

**disorder_pop[24] = 0.0464**

**disorder_pop[17] = 0.0494**

**disorder_pop[23] = 0.0494**

**disorder_pop[18] = 0.0517**

**disorder_pop[22] = 0.0517**

**disorder_pop[19] = 0.0531**

**disorder_pop[21] = 0.0531**

**disorder_pop[20] = 0.0535**

{*** Reaction setup ***}

{initiate concentrations}

b_quad = A0 + B0 + (k_off / k_on)

**C0** = (b_quad - SQRT((b_quad * b_quad) - (4 * A0 * B0))) / 2

**INIT A[0..40] = (A0 - C0) * disorder_pop[i]**

INIT B = B0 - C**0**

**INIT C[0..40] = C0 * disorder_pop[i]**

INIT D[0..n**, 0..40**] = 0

**INIT final[0..40] = 0 {same as D[n], for use in ARRAYSUM()}**

{modulating k_step}

slowdown = 300

k_step**[0..40]** = k_step_init**[i]** * max(0, (1 - (slowdown * **ARRAYSUM(final[*])** / B)))

{*** Differential equations ***}

d/dt (A**[0..40]**) = (C**[i]** * k_off) - (A**[i]** * B * k_on) + (D[n-1**, i**] * k_step**[i]**)

d/dt (B) = (**ARRAYSUM(C[*])** * k_off) - (**ARRAYSUM(A[*])** * B * k_on)

d/dt (C**[0..40]**) = (A**[i]** * B * k_on) - (C**[i]** * k_off) - (C**[i]** * **k_step[i]**)

d/dt (D[0**, 0..40**]) = (C**[j]** * k_step**[j]**) - (D[0**, j**] * k_step**[j]**)

d/dt (D[1..n-1**, 0..40**]) = (D[i-1**, j**] * k_step**[j]**) - (D[i**, j**] * k_step**[j]**)

d/dt (D[n**, 0..40**]) = (D[n-1**, j**] * k_step**[j]**)

**d/dt (final[0..40]) = (D[n-1, i] * k_step[i])**

{*** Output ***}

signal = min(**ARRAYSUM(final[*])**, Lg) * brightness

**Supplementary Figure Legends**

**Fig. S1. The NanoLuc transport assay**

**a)** Raw transport data (for pSpy_4x_ DDDL, see below) in dark blue, with a single exponential fit to the background (before addition of ATP) as an orange dotted line. Subtracting background from raw data yields the transport signal (light blue). All transport reactions curves shown in the paper have background subtracted, and signal immediately following ATP addition adjusted to 0 where needed.

**b)** Transport of 1 µM pSpy-Pep86 into PLs, measured in a cuvette [Pereira et al]. A standard transport assay is shown in black. When 20 µM 'cold' pSpy (i.e. without Pep86) is added together with the ATP – which should inhibit reinitiation, and thus approximate single turnover conditions – the orange trace is produced. Adding 20 µM cold pSpy together with the pSpy-Pep86, meanwhile – which competes directly for transport sites – gives the teal trace.

**c)** The data in panel **b** normalised to maximum amplitude for each trace. Because amplitudes from transport reactions not performed directly in parallel are highly variable, this is a more reliable way to compare the data. The normal (black) and single turnover (orange) traces are very similar, suggesting that transport is largely single turnover under these conditions.

**d)** Representative transport traces when the concentration of pSpy-Pep86 is varied: red = 4 µM, orange = 2 µM, yellow = 1 µM, green = 0.5 µM, teal = 0.25 µM, blue = 125 nM, purple = 63 nM, magenta = 31 nM. Black lines are best fits to the single exponential + lag.

**e)** Data in panel **d**, normalised by amplitude from a single exponential + lag fit.

**Fig. S2. Complete sequences and construction of the pSpy_4x_ series.**

See methods for details

**Fig. S3. Control experiments with the pSpy_4x_ series**

**a)** Transport signal from 8 µM pSpy-Pep86 (black line), or a dilution series of pSpy with scrambled Pep86 (see Fig. 2a): 8 µM (red), 4 µM (orange), 2 µM (yellow), 1 µM (green), 0.5 µM (teal), 250 nM (blue) or 125 nM (blue). The tiny signal produced by scrambled pSpy-Pep86 (see inset) confirms that it is still able to be transported.

**b-e)** Solution binding of the pSpy_4x_ series to 200 pM 11S, at 4 µM (red), 2 µM (orange), 1 µM (yellow), 0.5 µM (green), 250 nM (teal), 125 nM (blue) and 62.5 nM (purple).

**f)** Lag as a function of internal 11S concentration for the Spy_4x_ series, with the average value for each shown as a straight line.

**g)** λ as a function of internal 11S concentration for the Spy_4x_ series.

**h)** λ as a function of [ATP] for the Spy_4x_ series. Lines are global fits to a weak binding equation.

**Fig. S4. Modelling a single turnover reaction**

If SecYEG no longer replenishes after transport is complete, it causes transport to become susceptible to changes in SecYEG or 11S concentration (panel **b**), but not n, *k*_step_ or [pSpy-Pep86] (panels **c**-**e**). Therefore, this effect can not be responsible for controlling the signal amplitude.

**a)** Schematic of the model in section S2.2, in the same style as Fig. 1b. Once transport is complete, SecYEG is unable to mediate further rounds of pre-protein transport (represented by red X)

**b)** Effect of varying [SecYEG] on model in panel **a**.

**c-e)** Effect of varying **e)** n, **f)** *k*_step_ or **g)** [pSpy-Pep86].

The varied values are shown in the individual panels; default values are n = 5, *k*_step_ = 5, [pSpy-Pep86] = 1 µM, [11S] = 5 nM, *k*_on_ = 1.25 and *k*_off_ = 0.75. [SecYEG] is 8.4 nM in panels **a-c** and 4.2 nM in panels **e-g**.

**Fig. S5. Modelling SecYEG depletion over time**

**a)** Schematic of the model in section S2.3, in the same style as Fig. 1b. Here, we add an additional rate constant (*k*_Sec_death_) that depopulates all the translocation intermediates over time (red arrows).

**b-d)** Effect of varying n on model S2.3 with k_Sec_death set to **a)** 0, **b)** 1, or **c)** 2. If *k*_Sec_death_ is set high enough to slow transport appreciably, it makes the signal amplitude hypersensitive to n – far more than is observed experimentally.

**Fig. S6. Modelling transport with slowdown.**

**a)** Spy_4x_ series simulated using the model in S2.5; parameters are n = 5, *k*_step,init_ = 5, [pSpy-Pep86] = 1 µM, [SecYEG] = 1 nM, [11S] = 2.5 nM, *k*_on_ = 1.25 and *k*_off_ = 0.75, slowdown = 3000, brightness = 987. With these parameters, signal amplitude goes down as the position of the active Pep86 moves later – just as it does in the experimental data (Fig. 2b).

**b)** Effect of varying [pSpy-Pep86] on the transport curves produced by model S2.4. pSpy-Pep86 concentrations are listed in the figure; other parameters are n = 5, *k*_step_ = 5, [SecYEG] = 8.4 nM, [11S] = 5 nM, *k*_on_ = 1.25, *k*_off_ = 0.75, slowdown = 300, brightness = 200. Here, signal amplitude responds to pSpy-Pep86 concentration as long as [pSpy-Pep86]_in_ < [11S] when the reaction ends – just as it does experimentally (Fig. 1d).

**c)** Effect of varying *k*_step_ on the transport curves produced by model S2.4. Values for *k*_step_ are listed in the figure; other parameters are n = 5, [pSpy-Pep86] = 1 µM, [SecYEG] = 1 nM, [11S] = 2.5 nM, *k*_on_ = 1.25, *k*_off_ = 0.75, slowdown = 3000, brightness = 3000.

**Fig. S7. Effect of pSpy concentration on signal amplitude**

Normalised amplitudes from the individual pSpy titrations that comprise the averaged data in Fig. 1d. Each one is fitted to a straight line (where transport is limited by pSpy-Pep86 concentration) that caps at the point where the internal concentration of 11S is equal to the internal concentration of pSpy-Pep86. Further import has no effect on signal. Note the huge variation in intercept point (representing SecA motor strength – see main text), which arises because this value is susceptible to error from many sources, including the concentrations of 11S, pSpy-Pep86 and furimazine, instrument response, and the size distribution of PLs (see Fig. S8). Of these, [11S] is a particular problem: we add 40 µM 11S to reaction buffers when the PLs are prepared, then assume that this is evenly distributed inside and outside the PLs and does not change through the washing steps. To use signal amplitude as a measure of relative motor strength it is therefore crititally important to perform experiments with the same PLs, ideally in parallel using the same master mix.

**Fig. S8. Vesicle size distribution affects apparent kinetics**

**(a-c)** Small PLs **(a)** have a high surface:volume (and thus SecYEG:11S) ratio, meaning they fill up rapidly. Large PLs **(c)**, meanwhile, take much longer for tranport to reach completion – either by 11S running out (illustrated) or the concentration building up inside.

**d)** Simulated transport curves (model S2.1; red, orange, yellow, green, teal) with identical parameters (n = 20, *k*_step_ = 20), but containing 5 different concentrations of 11S. Brightness is scaled so each NanoLuc gives the same amount of signal. Adding all five traces together then normalising to 1 gives the black dotted line. The beginning of the curve looks very similar to the individual traces, but the slope phase is steeper, and the end tapers off in a completely different manner to the individual traces.

**e)** Measured the size distribution of PLs (extruded to 400 nm, exactly like those used in transport) by Dynamic Light Scattering (DLS; black line). The estimated surface area and volume of each vesicle subpopulation is calculated from 4πr^2^ and ^4^/_3_πr^3^, respectively (making the assumption that the PLs are spheres), then multiplied by population to give its share of the total surface area and volume (magenta and cyan, respectively). Small vesicles account for most of the surface area (magenta), but because volume scales with the cube of radius, the volume distribution skews towards the small number of large PLs (cyan).

**f)** To account for vesicle size distribution in Berkeley Madonna we modified model S2.4 to simulate each vesicle population separately, assuming that SecYEG is distributed uniformly across the vesicle surfaces and 11S is spread evenly through the encapsulated volume (S2.6). Simulated transport curves (with slowdown = 0) either without (teal line; S2.4) or with (red line; S2.6) vesicle size distribution modelled show the same effect as in panel **d**. Other parameters are identical: n = 5, *k*_step,init_ = 5 min^-1^, [pSpy-Pep86] = 1 µM, [SecYEG] = 8.4 nM, [11S] = 5 nM, *k*_on_ = 1.25 min^-1^, *k*_off_ = 0.75 min^-1^ and brightness = 200. The dotted line shows the best fit (by eye) to the red line to model S2.4; parameters are the same, except n = 6 and *k*_step,init_ = 7. The size distribution effect therefore potentially leads us to slightly overestimate both the number of steps and their rate

**g)** Same as panel **f**, but with slowdown set to 200 (for teal line) or 2000 (for red line), to give the same amplitude. Dotted line has the same parameters except slowdown = 500, *k*_step,init_ = 4.4 min^-1^ and brightness = 505, meaning the value for slowdown is very susceptible to population distribution.

**Fig. S9. The effect of static disorder on transport**

**(a)** To model static disorder in Berkeley Madonna we used a similar method as for vesicle fullness (Fig. S8f-g): the SecYEGs were divided into a set of different populations (41; 11 illustrated) populated according to a Gaussian distribution. These were and assigned a range of different intrinsic *k*_step_ rates (from *k*_step_/σ to *k*_step_×σ, where σ is a variable disorder parameter.

**(b)** Effect of including disorder as a parameter in the basic transport model. In general higher values for σ shorten the lag and slow the later stages of transport, as expected. Although this model is relatively cumbersome to work with, and produces complicated responses to changes in the different parameters, it does confirm that the number of steps determined from fitting should be considered a lower bound, as for other motor mechanisms.

**Fig. S10. ATP turnover rate of pSpy**

ATP hydrolysis rate as a function of pSpy concentration, showing average and SEM of three repeats. Fit is to the Michaelis Menten equation (S1.2), giving *k*_cat_ = 13.8 ± 1.2 s^-1^ and K_m_ = 0.6 ± 0.2 µM.
