## Supplementary figures for "Refined measurement of SecA-driven protein transport reveals indirect coupling to ATP turnover"

### Figure S1 – The NanoLuc transport assay

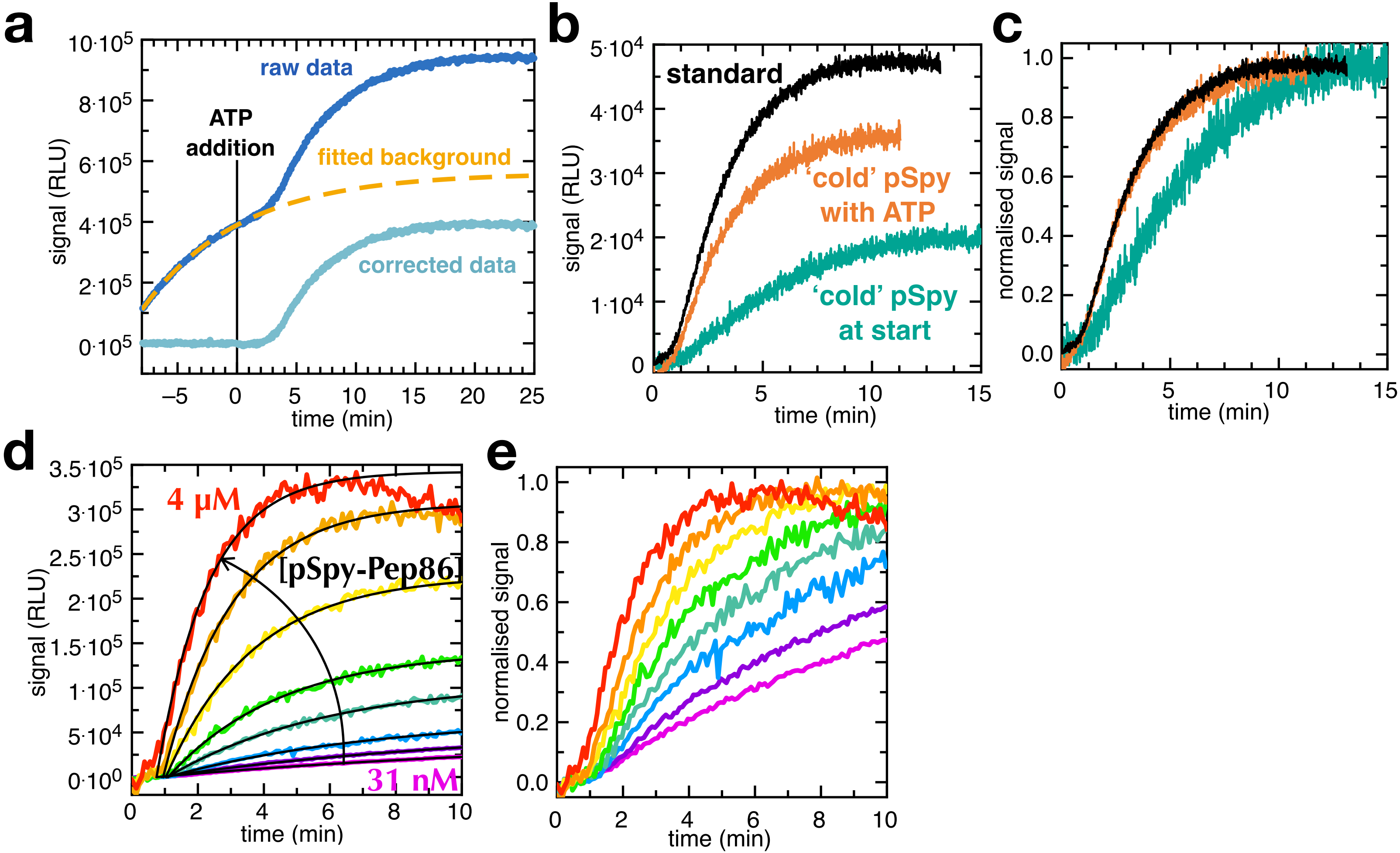

### Fig. S2 – pSpy tandem series sequences and cloning procedure

VSGWRLFKKIS = 'light' pep86
VSWGLRKFKIS = 'dark' pep86
MRKLTALFVASTLALGAANLAHA = signal sequence
HHHHHH = His tag
ENLYFQG = TEV site
IPNPLLGL = V5 epitope
DV = ZraI site

#### 1 - Synthesised Spy DD, DL and LD fragments

ADTTTAAPADAKPMMHHKGKFGPHQDMMFKDLNLTDAQKQQIREIMKGQRDQMKRPPLEERRAMHDIIASDTFDKVKAEAQIAKMEEQRKANML
AHMETQNKIYNILTPEQKKQFNANFEKRLTERPAAKGKMPATAEGSGXXXXXXXXXXXGSGADTTTAAPADAKPMMHHKGKFGPHQDMMFKDLN
LTDAQKQQIREIMKGQRDQMKRPPLEERRAMHDIIASDTFDKVKAEAQIAKMEEQRKANMLAHMETQNKIYNILTPEQKKQFNANFEKRLTERP
AAKGKMPATAEGSGXXXXXXXXXXXGSG

#### 2 - Previously published pBAD-pSpy-V5-pep86-TEV-His sequence

MRKLTALFVASTLALGAANLAHAADTTTAAPADAKPMMHHKGKFGPHQDMMFKDLNLTDAQKQQIREIMKGQRDQMKRPPLEERRAMHDIIASD
TFDKVKAEAQIAKMEEQRKANMLAHMETQNKIYNILTPEQKKQFNANFEKRLTERPAAKGKMPATAEIPNPLLGLGSGVSGWRLFKKISENLYF
QGHHHHHH

DD, DL and LD fragments cloned into pBAD -V5-HiBiT-TEV-His in place of Spy-V5. Primers include ZraI site.

#### 3 - pBAD-pSpy(DD, DL and LD)-ZraI-TEV-His

MRKLTALFVASTLALGAANLAHAADTTTAAPADAKPMMHHKGKFGPHQDMMFKDLNLTDAQKQQIREIMKGQRDQMKRPPLEERRAMHDIIASD
TFDKVKAEAQIAKMEEQRKANMLAHMETQNKIYNILTPEQKKQFNANFEKRLTERPAAKGKMPATAEGSGXXXXXXXXXXXGSGADTTTAAPAD
AKPMMHHKGKFGPHQDMMFKDLNLTDAQKQQIREIMKGQRDQMKRPPLEERRAMHDIIASDTFDKVKAEAQIAKMEEQRKANMLAHMETQNKIY
NILTPEQKKQFNANFEKRLTERPAAKGKMPATAEGSGXXXXXXXXXXXGSGDVGSGENLYFQGHHHHHH

Synthesised DD, DL and LD fragments amplified by linear PCR and cloned into ZraI site

#### 4 - pBAD-pSpy(DDDD, LDDD, DLDD, DDLD, DDDL)-ZraI-TEV-His

MRKLTALFVASTLALGAANLAHAADTTTAAPADAKPMMHHKGKFGPHQDMMFKDLNLTDAQKQQIREIMKGQRDQMKRPPLEERRAMHDIIASD
TFDKVKAEAQIAKMEEQRKANMLAHMETQNKIYNILTPEQKKQFNANFEKRLTERPAAKGKMPATAEGSGXXXXXXXXXXXGSGADTTTAAPAD
AKPMMHHKGKFGPHQDMMFKDLNLTDAQKQQIREIMKGQRDQMKRPPLEERRAMHDIIASDTFDKVKAEAQIAKMEEQRKANMLAHMETQNKIY
NILTPEQKKQFNANFEKRLTERPAAKGKMPATAEGSGXXXXXXXXXXXGSGDADTTTAAPADAKPMMHHKGKFGPHQDMMFKDLNLTDAQKQQI
REIMKGQRDQMKRPPLEERRAMHDIIASDTFDKVKAEAQIAKMEEQRKANMLAHMETQNKIYNILTPEQKKQFNANFEKRLTERPAAKGKMPAT
AEGSGXXXXXXXXXXXGSGADTTTAAPADAKPMMHHKGKFGPHQDMMFKDLNLTDAQKQQIREIMKGQRDQMKRPPLEERRAMHDIIASDTFDK
VKAEAQIAKMEEQRKANMLAHMETQNKIYNILTPEQKKQFNANFEKRLTERPAAKGKMPATAEGSGXXXXXXXXXXXGSGVGSGENLYFQGHHH
HHH

### Fig. S3 – 4x tandem spies supporting data

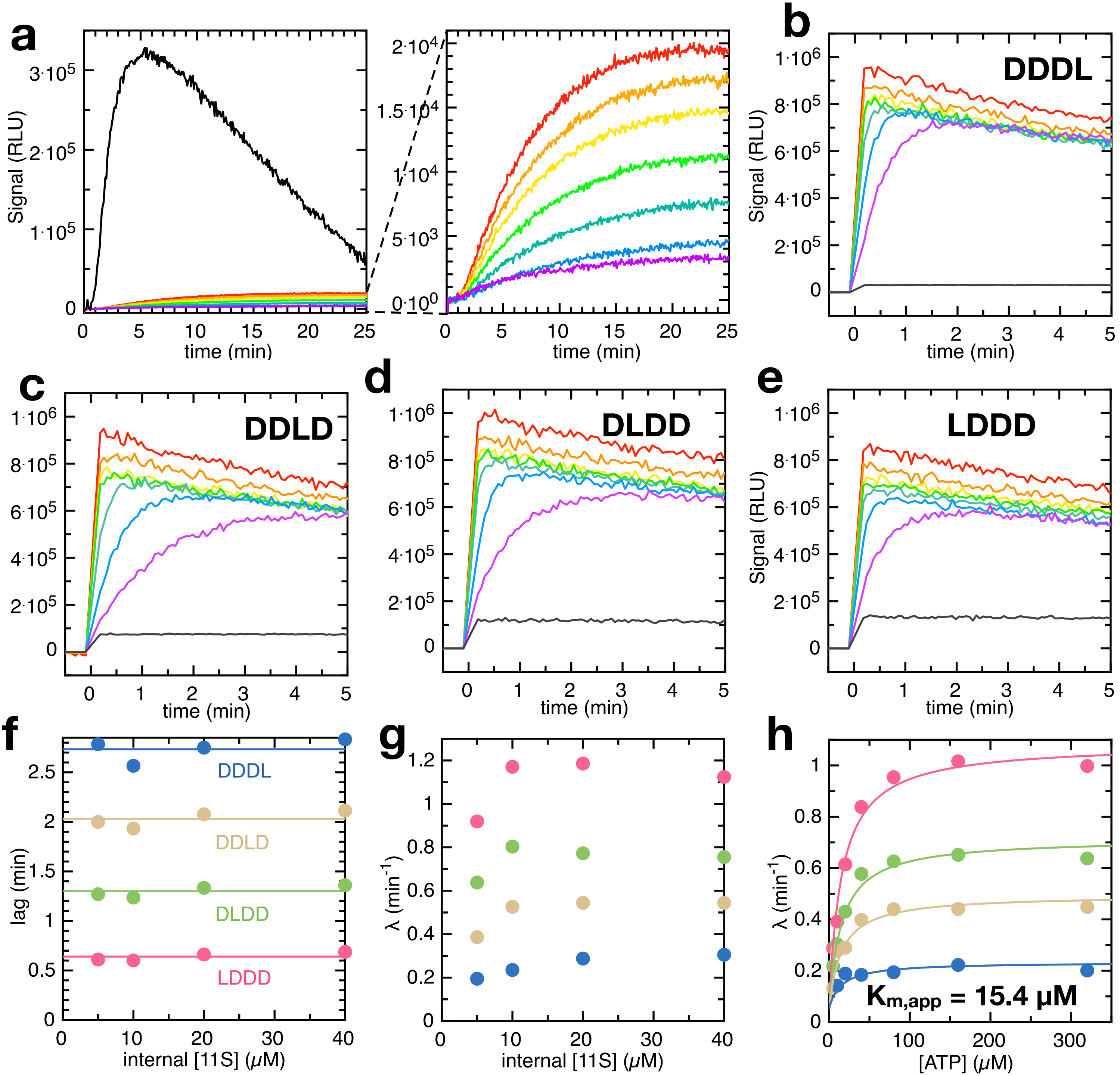

### Fig. S4 – Modelling a single turnover transport reaction

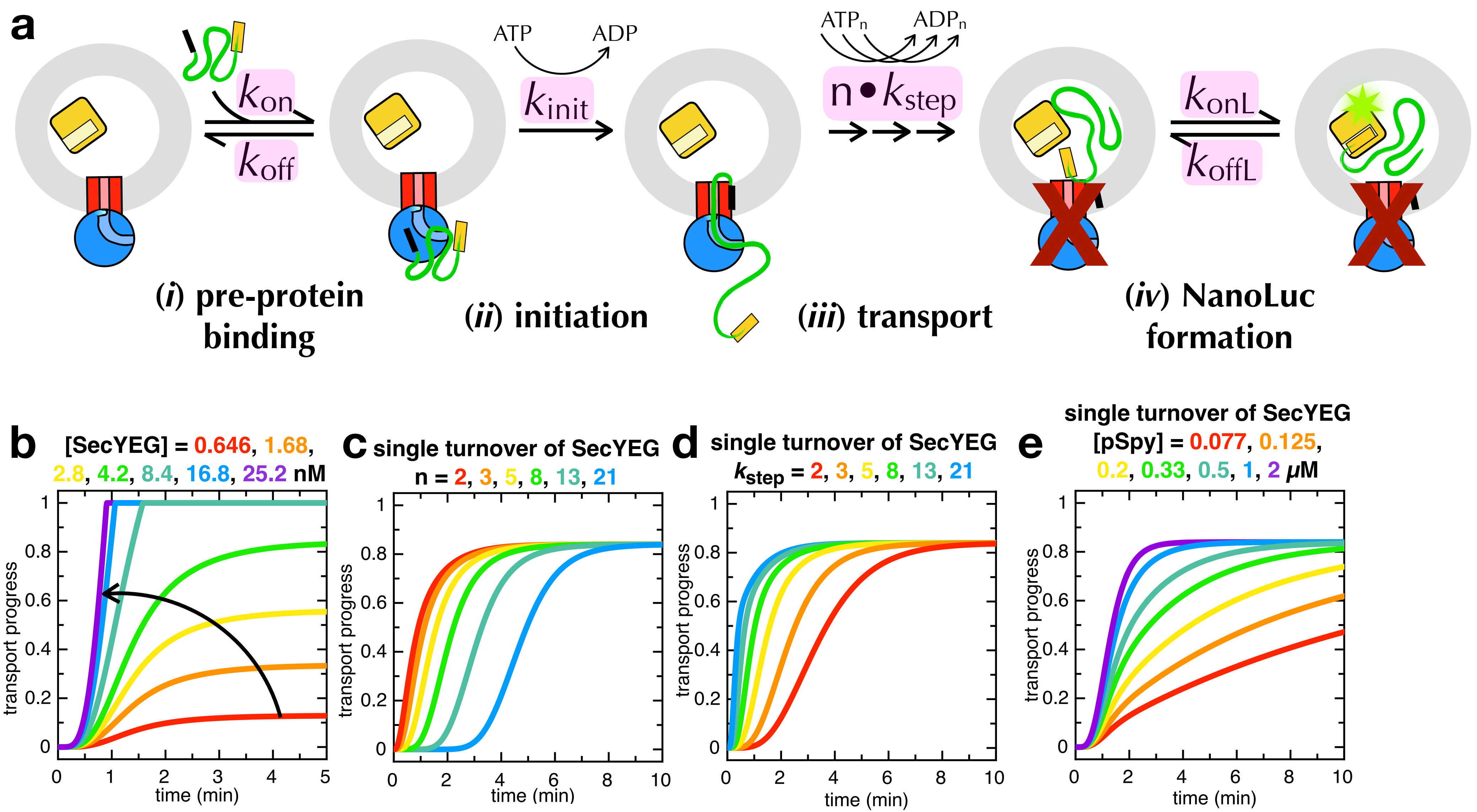

### Fig. S5 – Modelling SecYEG depletion over time

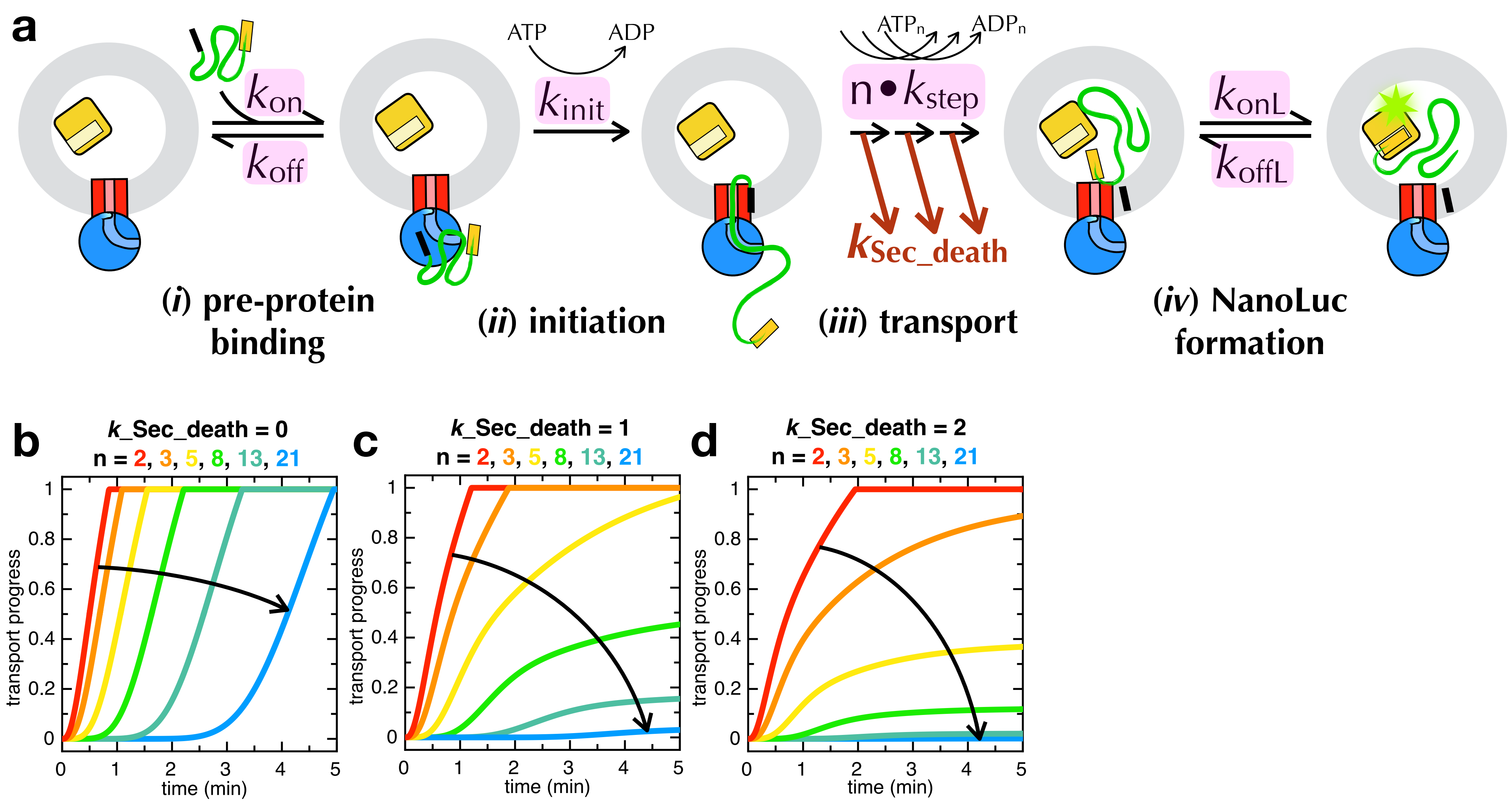

### Fig. S6 – Modelling transport with slowdown

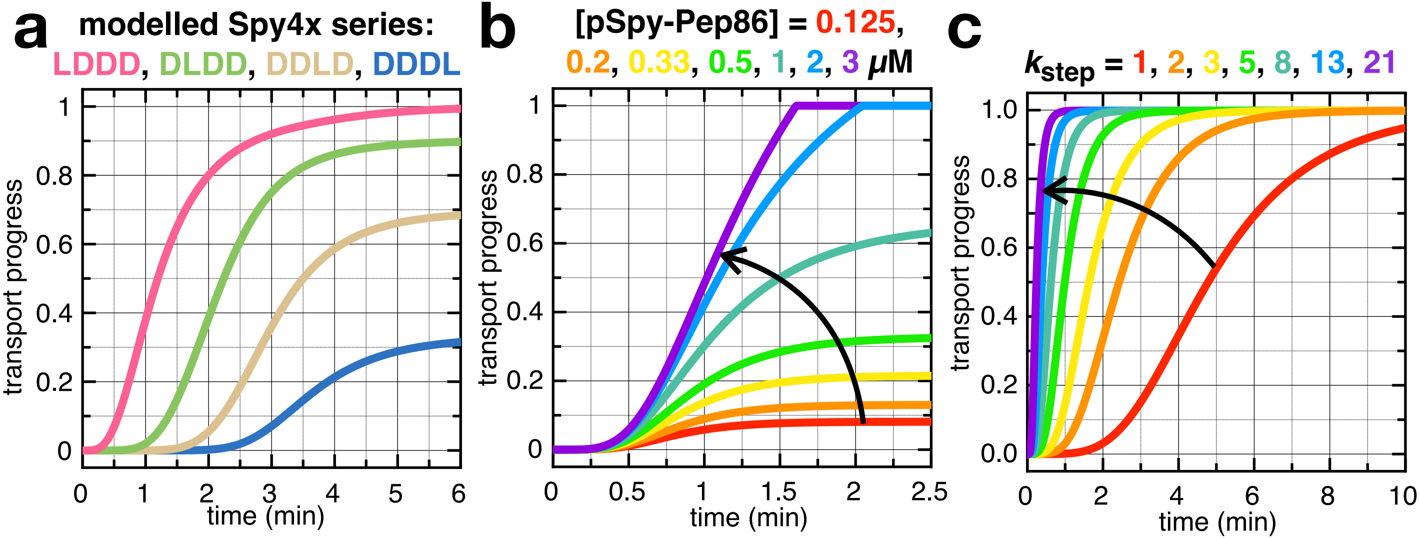

### Fig. S7 – [pSpy] and amplitude

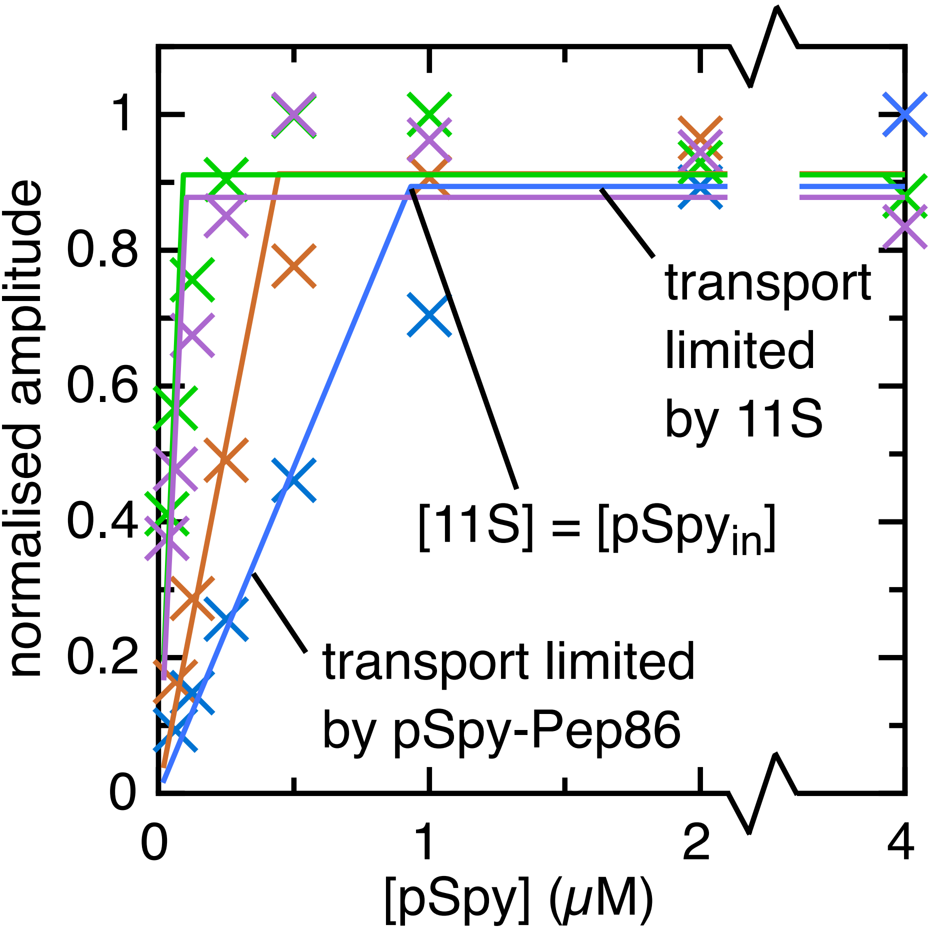

### Fig. S8 – surface/volume correction

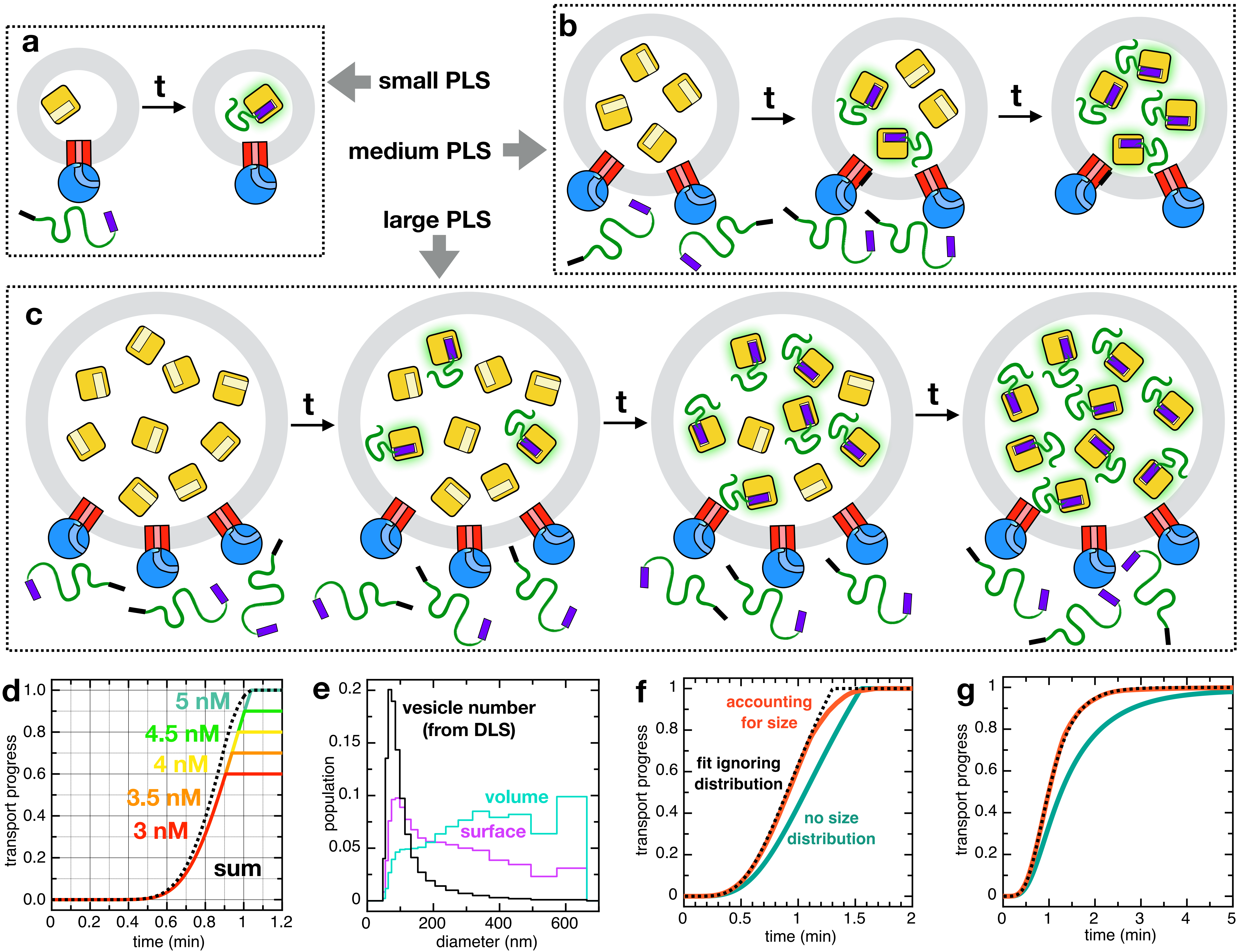

### Fig. S9 – Static disorder

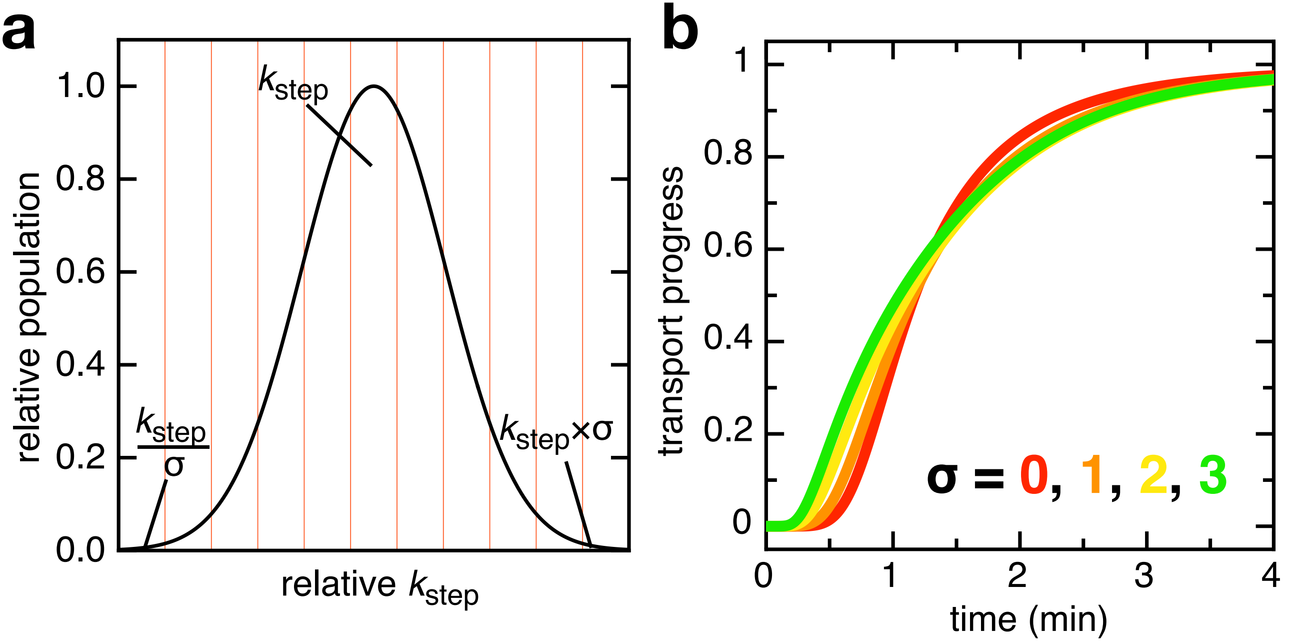

### Fig. S10 – SecA ATPase activity in pSpy transport

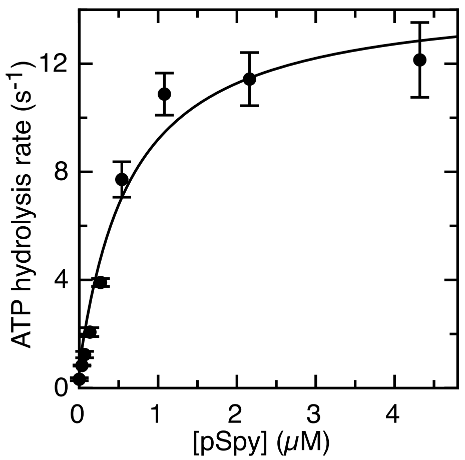
